# With a pinch of salt: metagenomic insights into Namib Desert salt pan microbial mats and halites reveal functionally adapted and competitive communities

**DOI:** 10.1101/2022.02.18.481119

**Authors:** Laura Martínez-Alvarez, Jean-Baptiste Ramond, Surendra Vikram, Carlos León-Sobrino, Gillian Maggs-Kölling, Don A. Cowan

**Affiliations:** Centre for Microbial Ecology and Genomics (CMEG), Department of Biochemistry, Genetics and Microbiology, University of Pretoria, Pretoria, South Africa; Departamento de Genética Molecular y Microbiología – Pontificia Universidad Católica de Chile – Chile; Gobabeb-Namib Research Institute, Walvis Bay, Namibia

**Keywords:** Salt pan, playa, CRISPR-Cas, functional diversity, taxonomic diversity, microbial mat, halite, horizontal gene transfer, Gene Transfer Agent, Virus-Host interactions

## Abstract

Salt pans or playas, which are saline-rich springs surrounded by halite evaporates in arid environments, have played an essential role in landscape erosion during the formation of the Namib Desert and are numerous in its central region. In this study, we used shotgun metagenomics to investigate the phylogenetic and functional capacities of the microbial communities from two salt pans (namely, Eisefeld and Hosabes) located in the Central Namib Desert, located in Southwest Africa. We studied the source and sink sediment mat communities of the saline streams, as well as those from two halites (crystallized structures on the stream margins). The microbial assemblages and potential functions were distinct in both niches. Independently from their localization (Eisfeld vs Hosabes and source vs sink), the sediment mat communities were dominated by members of the *Alpha-* and *Gamma-proteobacteria* classes, while halites were Archaea-dominated and also contained high abundances of the extremely halophilic bacterium *Salinibacter* sp. (phylum *Bacteroidota*). Photoheterotrophy and chemoheterotrophy were the principal lifestyles in both niches, with halite communities having a reduced diversity of metabolic pathways. Intense microbial-virus interactions in both niches were implied by the widespread detection of CRISPR-Cas defense systems. We identified a putatively novel clade of type II CRISPR-Cas systems, as well as novel candidate viral lineages of the class Caudoviricetes and of Halobacteriales*-*infecting haloviruses. Putative gene transfer agent-like sequences within the *Alphaproteobacteria* were identified in the sediment mat communities. These horizontal gene transfer elements have the potential to drive genome plasticity and evolution of the *Alphaproteobacteria* in the Namib Desert salt pan microbiomes.

**Importance:** The hyperarid Namib Desert is one of the oldest deserts on Earth. It contains multiple clusters of playas which are saline-rich springs surrounded by halite evaporites. Playas are of great ecological importance and their indigenous (poly)extremophilic microorganisms are potentially involved in the precipitation of minerals such as carbonates and sulfates and have been of great biotechnological importance. While there has been a considerable amount of microbial ecology research preformed on various Namib Desert edaphic microbiomes, little is known about the microbial communities inhabiting its multiple playas. In this work, we therefore provide a comprehensive taxonomic and functional potential characterization of the microbial, including viral, communities of sediment mats and halites from two distant Namib Desert, contributing towards a better understanding of the ecology of this biome.

## Introduction

Saline inland waters account for 5% of dryland surfaces globally (1) and represent approximately 0.008% of the world’s water. This is almost equivalent to the total amount of freshwater, estimated to be of 0.009% (2). Salt pans - or playas - are terrain depressions found in arid ecosystems where underground water surfaces (source) and evaporates along its stream - which leads to the formation of a salt-crust over the ground sediment - before disappearing again underground at its sink (3). The Namib Desert is one of the oldest deserts in the world, estimated to have been hyper-arid for the last 5 million years (4), and is characterized by the presence of numerous playas (3). Although salt pans occupy less than 5% of the central Namib Desert gravel plains, they are a major water source for the desert fauna and play an important role in the Namib Desert’s geomorphology via gypsum (CaSO_4_*2H_2_0) deposition and landscape erosion through salt weathering (3, 5, 6). Furthermore, they produce some of the most saline inland waters in southern Africa, with measured salinities reaching up to 160 grams of total dissolved solids per liter (values >3 g/L are considered saline water) (5, 7).

The microbial diversity of salt pans worldwide is dependent on their particular geochemical characteristics, including salinity, pH and oxygen levels. In salt pans, microbial communities develop into microbial mats with a vertical-layered structure in which each layer harbors different microorganisms with distinct metabolic capacities (8). Globally, microbial diversity decreases with increasing salinity, and it is accompanied by increasing proportions of Archaea; particularly of the order *Halobacteriales* from the *Halobacteriota* phylum (9–11). Furthermore, the taxonomic diversity of saline microbial mat communities fluctuates in response to the constant changes in their surrounding physicochemical conditions; particularly in response to salinity, oxygen levels and their metabolic activity (12). While many studies have addressed the taxonomic diversity of saline microbial mats around the world (9–11), their functional capacities remain largely unexplored. Similarly, while hypersaline environments are known to be rich in novel viruses (e.g., (13–16)) that are likely to modulate microbial community compositions and biogeochemical cycling functions (17), their ecological roles have largely been uninvestigated. Previous studies confirmed that Namib Desert saline environments and show a wealth of novel viruses (13). However, little information on virus-host pairs in such environments is available.

Consequently, in order to investigate the taxonomic composition and functional potentials of microbial and viral communities in desert playa microbial mats and associated salt crusts (i.e., halites), we investigated ten shotgun metagenomes from two central Namib Desert playas belonging to two different saline spring clusters (3, 6, 7). We noted that the microbial and viral mat communities from both distant salt pans were highly similar in their taxonomic distribution and functional potential. Conversely, halites were dominated by taxa which are typically adapted to hypersaline conditions and low water availability, with relatively lower phylogenetic diversity. Novel viral taxa dominated both mat and halite communities, with a higher phylogenetic diversity in the former being consistent with the greater microbial diversity of mat microbiomes. Analyses of the defense systems used by the community against mobile genetic elements revealed abundant type I and type III CRISPR-Cas systems, as well as putative novel subtype of type II systems. Additionally, a cluster of gene transfer agents (GTAs) with the potential to mediate horizontal gene transfer events was identified in the mat communities. Overall, our results strongly suggest that the saline spring microbial mat and halite niches represent microbial and viral diversity hotspots, characterized by diverse functional capacities, high inter-taxon competition and high capacities for genomic evolution and adaptation.

## Results

Eight metagenomes were generated from the microbial salt pan mats of the Namib Desert Hosabes and Eisfeld playas (Figure 1). The samples were collected from their source and sink of the water stream in 2016 and 2017. Additionally, two more metagenomes were produced from crystalline halites collected in the vicinity of the Hosabes playa stream; named as “dark”- and “red”-halite due to the surface color of the rock (Figure 1). The 10 metagenomes comprised between 1.04 (Hosabes source 2016) and 9.57 (Red halite) Gbp of sequencing data. After data processing and assembly, 4 527 165 contigs over 500 bp were retained for further analyses (Supplementary Table 1). Nonpareil analyses clearly showed that that the sequence depth was high, covering between 66% (Eisfeld source 2017) to 90 % (Hosabes sink 2017) of the microbial communities in all samples (Supplementary Figure 1).

**Figure 1.**
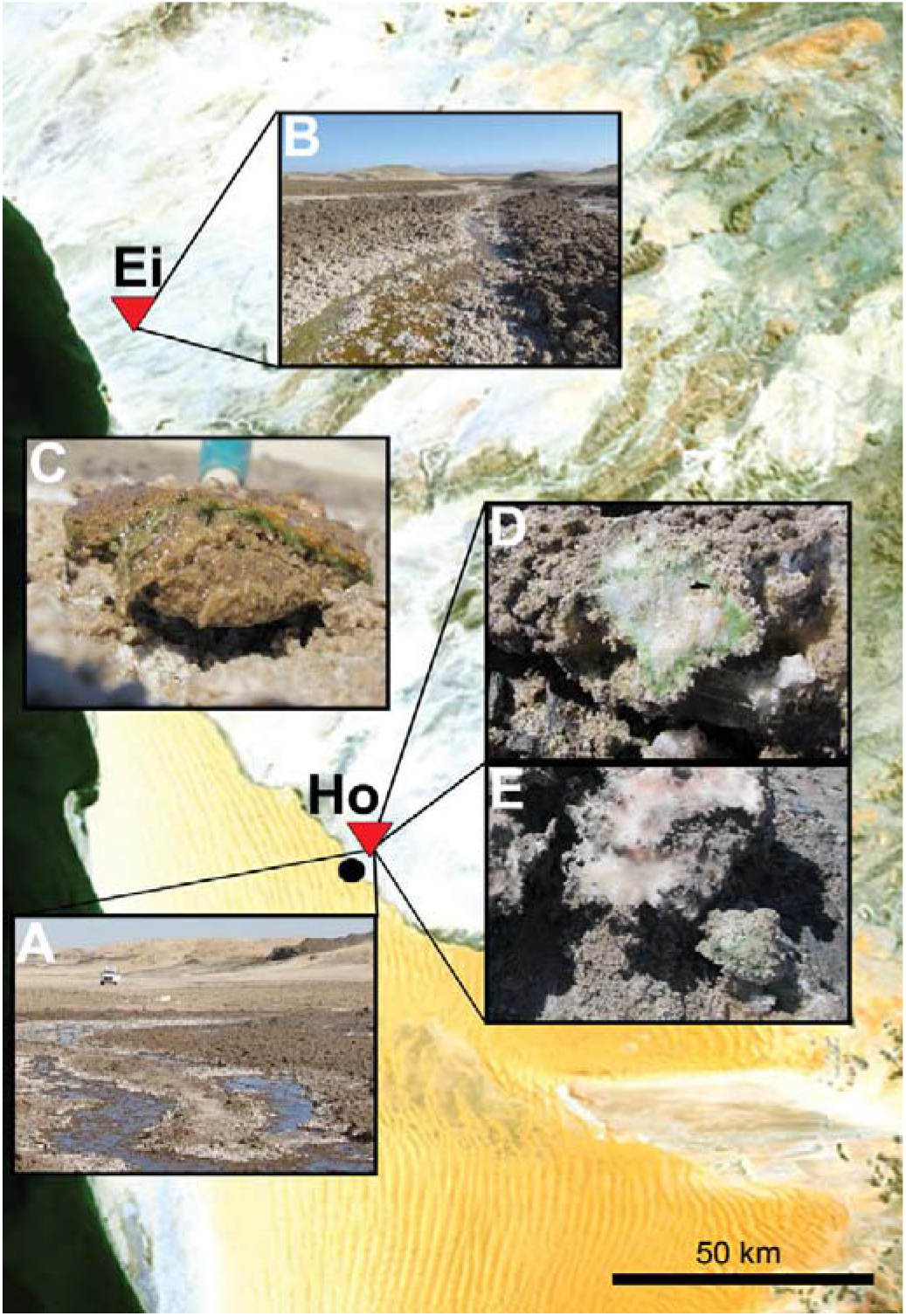
Map of the central Namib Desert with the sampling location distribution. It clearly shows the extensive northern gravel plains and the southern dune fields. The red triangles indicate the location of the two sampled playas - Hosabes (Ho) and Eisfeld (Ei) - and the black dot the Gobabeb – Namib Research Institute. Inset photographs depict the salt pan streams of Hosabes **(A)** and Eisfeld **(B)**, and a close-up of the collected samples: microbial mat **(C)**, and the dark **(D)** and red **(E)** halite rocks. Image adapted from ESA, CC BY-SA 3.0 IGO.

### Niche-specific microbial assemblages in Namib Desert hypersaline environments

Around 65.5% of the coding sequences predicted from the metagenomic data could taxonomically be assigned at Order level (Supplementary Table 1) and were used to profile microbial community diversity. Halite and mat metagenomes displayed significantly different taxonomic composition as shown by their clear separation on the PCA plot of Figure 2A. Furthermore, both halite communities were found more dissimilar than those of the mats (independently from year and sample type [source vs sink]). In contrast, the mat communities of both playas were rather similar as the variance between the Eisfeld and Hosabes mat metagenomes was comparable to the variance between their source and sink communities (Figure 2A), with an average 5% difference in taxonomic diversity.

**Fig 2.**
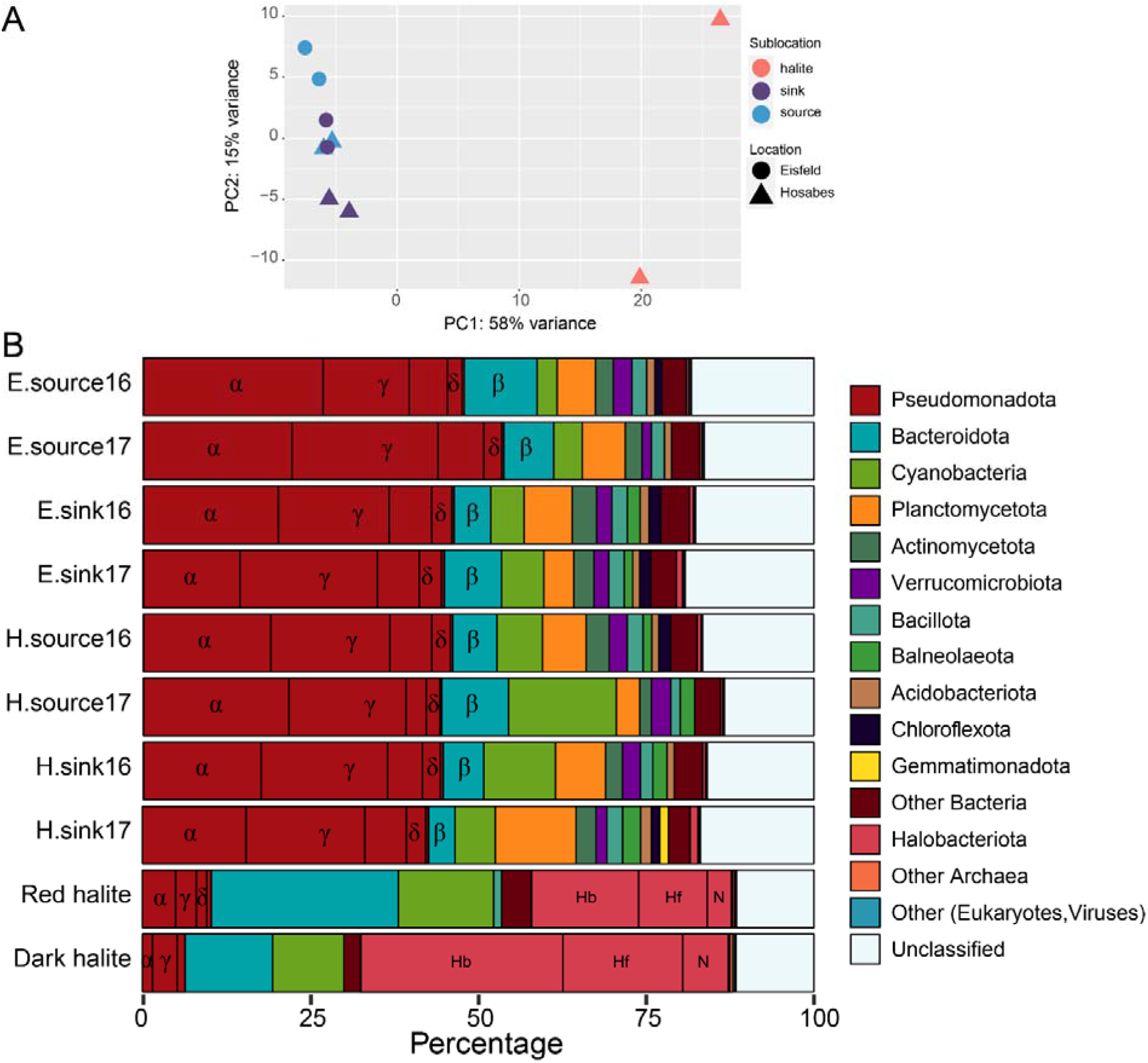
Namib Desert microbial community mat and halite samples diversity. **A)** Principal component plot of the Namib stream mat and halite samples based on gene taxonomy at Order level obtained using the DESeq2 package. Taxonomic diversity was compared by location (e.g., Eisfeld or Hosabes) and sublocation (e.g., source, sink or halite). **B)** Relative taxonomic classification of the metagenomic open reading frames. **Proteobacteria** dominate the salt pan samples, while members of the *Euryarchaea* are predominant in both halites. H – Hosabes, E – Eisfeld, α – *Alphaproteobacteria*, β – *Betaproteobacteria*, γ – *Gammaproteobacteria*, δ – *Deltaproteobacteria*, Hb – Halobacteriaceae, Hf – Haloferacaceae, N – Natrialbaceae.

The halite microbial communities were clearly Archaea-dominated when compared to the stream mats (Figure 2B). *Halobacteriota* represented 55.4% of the dark halite and 30.3% of the red halite. The *Halobacteriales* order particularly dominated both archaeal fractions, with relative abundances ranging from 30.3% (red halite) to 55.1% (dark halite; Figure 2B and Supplementary Table 5). The halite bacterial fraction was also dominated by well-known salt-tolerant/halophilic genera - particularly *Salinibacter sp.* (*Bacteroidota* phylum)*, Halothece* sp. and *Dactylococcopsis sp*. (*Cyanobacteria*; Supplementary Table 5) - while these were not abundant in the saline spring mat assemblages. Furthermore, the halite bacterial fractions presented a 40-fold enrichment in *Rhodothermaceae* sp. (*Bacteroidota*)*, Aphanothecaceae* sp. (*Cyanobacteria*) and *Wenzhouxiangellaceae* sp. (*Gammaproteobacteria* class) in comparison to the mat assemblages (Table 1).

**Table 1.**
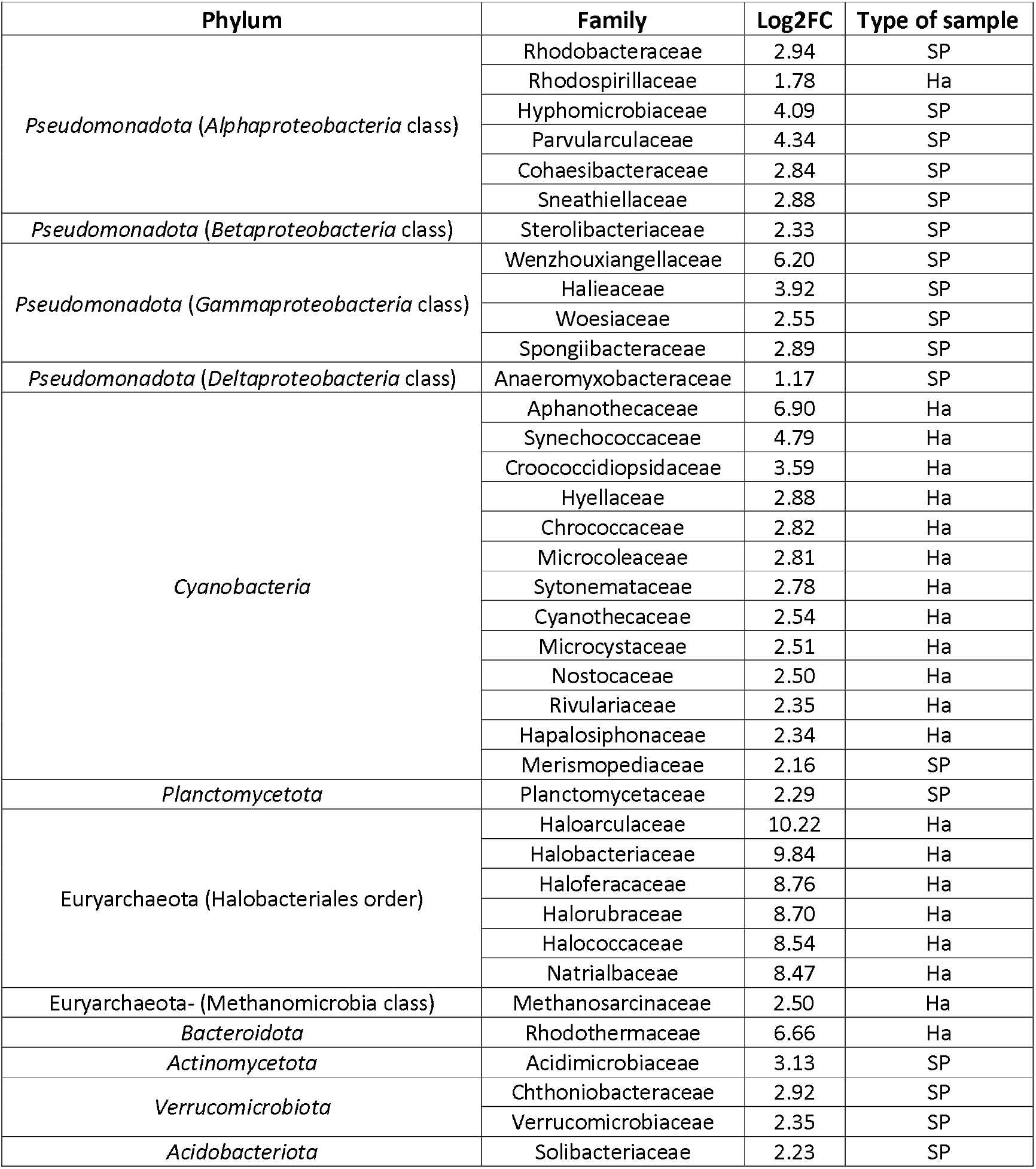
Principal differences in taxonomic composition of salt pan and halite samples at family level. Fold change differences (log2FC) in taxon abundance between stream mat and halite samples. The column “Type of sample” indicates in which community (SP: Salt Pan; Ha: Halite) is the taxon enriched.

Stream mat microbial communities comprised a total of 11 bacterial phyla and were dominated by members of the *Pseudomonadota* phylum, and particularly of the *alpha-* and *gamma-proteobacteria* classes with relative abundances ranging from 14.4 to 26.8% and 12.8 to 21.8% of the total mat communities, respectively (Figure 2B). The other dominant mat bacterial phyla were *Bacteroidota* (5.1-10%), *Cyanobacteria* (2-13%) and *Planctomycetota* (3.4-11.8%). *Actinomycetota*, *Verrucomicrobiota*, *Bacillota*, *Balneolota* and *Acidobacteriota* comprised 1% to 3.6% (Figure 2B). We noted that the cyanobacterial Aphanothecaceae and Leptolyngbyaceae and the alphaproteobacterial Methylocystaceae families were slightly enriched in Hosabes salt pan metagenomes (4-fold as calculated using DESeq2 from the normalized counts, see Methods), while the Eisfeld samples were enriched the cyanobacterial Coleofasciculaceae and alphaproteobacterial Hyphomicrobiaceae family members (3-4-fold) (Supplementary Table 3). Furthermore, members of the alphaproteobacterial Rhodobacteraceae, Methylocystaceae and Parvularculaceae families, and of the *Bacteroidota* Flavobacteriaceae family, were 2-3 fold more abundant in mat sources, while members of the *Nitrospinota, Hydrogenedentes* and *Balneolota* phyla were 2-3-fold more abundant in mat sink communities (Supplementary Table 4).

Altogether, our results clearly show that, despite belonging to 2 different saline stream “clusters” separated by over 120 km (Figure 1), the Eisfeld and Hosabes mat communities are highly similar (Figure 2). Conversely, those of the halite communities, which were only 50 m apart, were significantly different (Figure 2).

### The functional capacities of the Namib Desert saline communities differ in halite and salt pans stream mats

An average of 39% of the metagenomic open reading frames (ORFs) could be assigned to KEGG Ortholog (KO) terms (Supplementary Table 1). Of the genes with KO term annotation, around 35% belong to the metabolism category and predominantly to the amino acid (between 8.6 to 9%) and carbohydrate (10.4-11.5%) metabolism subcategories (Supplementary Figure 2). In the environmental information processing category, we particularly noted that genes from the membrane transport subcategory accounted for approximately 0.1% of the total ORFs (Supplementary Figure 2). Genes related to signal transduction and glycan biosynthesis were approx. 50% less abundant in the halite than in the mat communities (Supplementary Figure 2).

Transport of osmoprotectant solutes (glycine/betaine/proline transport [M00208], osmoprotectant and polyamine transport systems [M00209]) orthologs were widespread in the stream mat metagenomes and encoded in sequences belonging to the bacterial *Alpha*-, *Gamma*-, and *Delta-proteobacteria* classes and *Bacteroidota* and *Cyanobacteria* phyla as well as to the archaeal Haloferacales order (Supplementary Table 6). This suggests these organisms employ a “salt-out” strategy to balance osmotic stress and therefore can withstand a range of salinities (18). By contrast, the absence of these systems in the archaeal *Halobacteriota* and bacterial *Planctomycetota* phyla suggests that members from these taxa rather employ a “salt-in” strategy (18). As expected, transport systems for lipopolysaccharide and other capsular polysaccharides were 3-8 times more abundant in the mat metagenomes than in the halite ones (Supplementary Table 6).

The nutrient (C, N, S) biogeochemical cycling capacities were also very different in both niches (Figure 3): Anaerobic C fixation, nitrification, denitrification, dissimilatory nitrate reductions, dinitrogen fixation and dissimilatory sulfate reduction (a form of anaerobic respiration performed by chemoorganoheterotrophic microbes; (19)) were exclusively found in the mat metagenomes.

**Figure 3.**
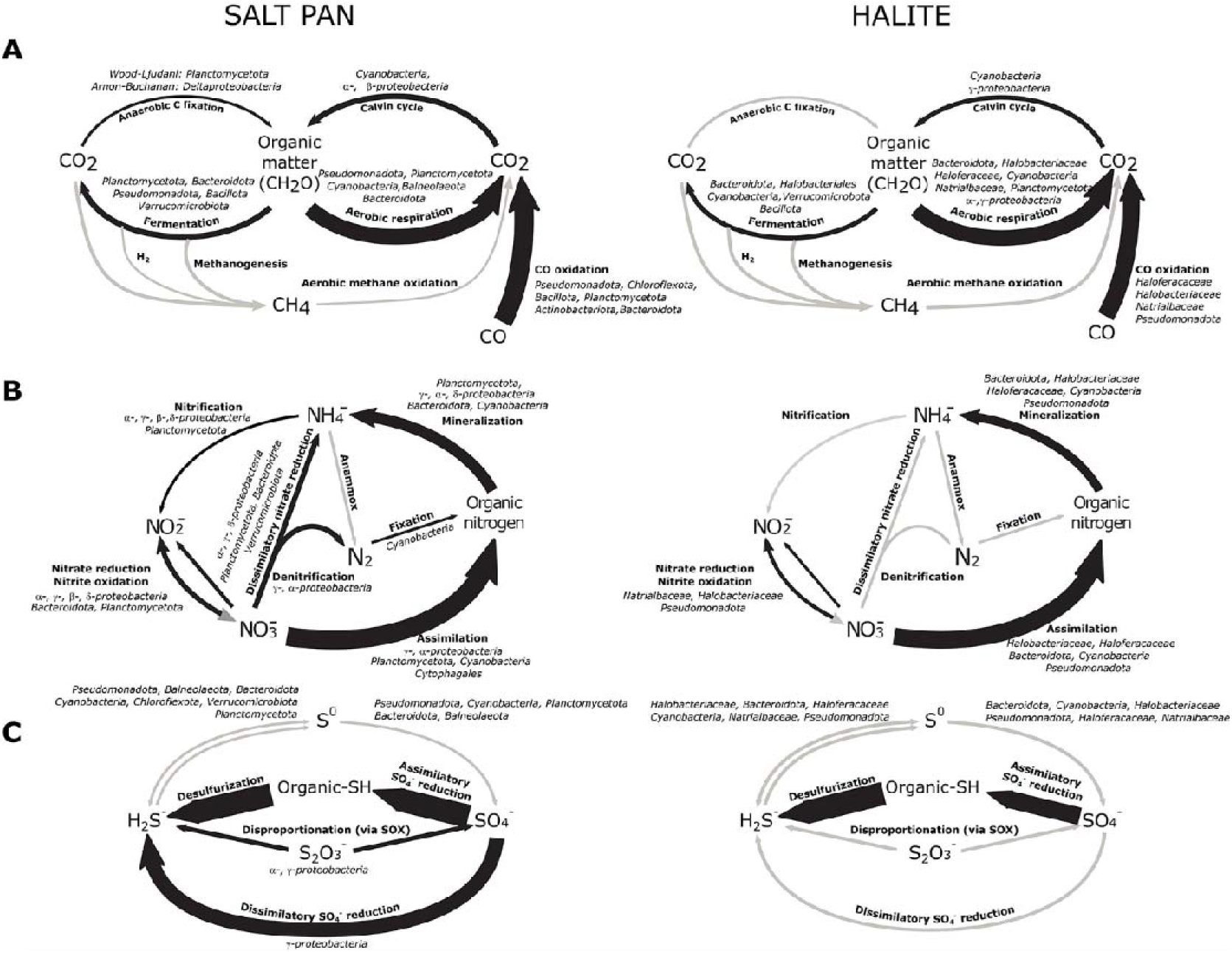
Functional potential of the Namib salt pan and halite microbial communities. Panels depict the carbon **(A)**, nitrogen **(B)** and sulfur **(C)** cycles from the mat (left) and halite (right) metagenomic data. Black arrows represent steps of the cycle present in the community, while grey arrows represent pathways not detected in the metagenomes. Thickness of arrows is proportional to the marker gene counts for each pathway. Main taxa with the genetic potential for each metabolic step are shown in italics.

Capacity for carbon fixation via the Calvin cycle was detected in the mat and halite communities and could be carried out by members of the *Cyanobacteria* and *Alpha*- and *Beta*-*proteobacteria*. Photosystem I and II modules were complete for the Cyanobacteria in niches, and this was the only phylum with capacity of oxygenic photosynthesis. Nevertheless, the capacity for phototrophy was widespread amongst the *Alphaproteobacteria,* where the anoxygenic photosystem II module was found complete in the mat metagenomes (Figure 3A and Supplementary Table 6). Members of the mat *Beta*- and *Gamma*-*proteobacteria* may also have the functional capacity for anoxygenic phototrophy since the *pufLM* genes coding for subunits of the photosynthetic reaction centre were found in these taxa (see Supplementary Table 7). Evidence for the presence of the anaerobic Arnon-Buchanan and Wood-Ljungdahl pathways of carbon fixation was restricted to the salt pan *Deltaproteobacteria* and *Planctomycetota*, respectively (Figure 3A). Additionally, the capacity to obtain energy from CO oxidation (carboxydovory) was widespread in both mat and halite communities, implying that the use of alternative energy sources may assist these microorganisms to survive under oligotrophic conditions. No evidence of the capacity for methanogenesis (i.e., lack of *mcr*A gene) was found in any of the metagenomes. Taken together, these data suggest that *Cyanobacteria* and *Alphaproteobacteria* as the predominant primary producers in the Namib Desert hypersaline ecosystems.

It is noted that alternatives to the TCA cycle, which is thought to be employed to avoid carbon loss (20), were widespread in all metagenomic datasets. The glyoxylate cycle was found in *Alpha*-, *Gamma*-*proteobacteria*, *Cyanobacteria*, *Bacteroidota* phyla and Halobacteriales order. The ethylmalonyl pathway was found exclusively in the salt pan *Alphaproteobacteria*, an expected finding given the high abundance of the family Rhodobacteraceae (21). The archaeal methylaspartate cycle, which has been described as an adaptation to halophilic conditions and carbon starvation (22), was only detected in the archaeal-dominated halite metagenomes (Supplementary Table 6), concurrent with the high presence of gene *phaC* [which encodes for for the polyhydroxyalkanoate (PHA) synthase] in the Halobacteriaceae and Haloferacaceae, as well as the *Alphaproteobacteria* (Supplementary Table 7). PHAs are major storage compounds in prokaryotes, and as the ethylmalonyl pathway is interrelated with the synthesis of PHAs, this cycle together with the methylaspartate pathway has been shown to be linked to the capacity to adapt to environmental stresses (23).

Only *Cyanobacteria* from the stream mat communities showed the potential capacity to fix atmospheric nitrogen, albeit that the genetic markers of this capacity (i.e., the *nif* genes) were present at low abundances. Assimilatory nitrate reduction capacity was found predominantly in the archaeal Halobacteriales and Haloferacales orders in halites, and in *Planctomycetota*, *Cyanobacteria* and *Alphaproteobacteria* in both saline systems (Figure 3B). This step was the only inorganic nitrogen incorporation reaction detected in the halite microbial community, suggesting an incomplete nitrogen cycle in halites. Complete modules for this pathway were detected in all main phyla *Alpha*-, *Gamma*-, *Delta*-*proteobacteria*, *Planctomycetota* and *Bacteroidota* (Figure 3B). Metabolic capacity for denitrification was found in the salt pan *Alpha*-, *Gamma-proteobacteria* and *Bacteroidota* (Figure 3B). Unlike dissimilatory nitrate reduction, denitrification does not conserve nitrogen in the system, which is lost as volatile nitrogen forms (N_2_). Some capacity for nitrification was detected for the salt pan *Alpha*-(Rhizobales and Rhodobacterales orders), *Beta*-(Nitrosomonadales order), *Gamma-proteobacteria* (Methycoccales, Chromatiales and Oceanospirillales orders) and *Planctomycetota*, (Supplementary Table 7). No evidence of anaerobic ammonium oxidation (annamox) was detected in either the salt pan or the halite samples.

Assimilatory sulfate reduction was the only sulfur incorporation step identified in the halite communities. Conversely, salt pan *Gammaproteobacteria* possessed the capacity for anaerobic respiration through the dissimilatory sulfate reduction pathway. Similarly, the presence of the genes for thiosulfate oxidation via the SOX complex suggested a capacity for chemolithoautotrophy for the salt pan *Alpha-* and *Gammaproteobacteria* (Figure 3C). These results suggest that the halite microbial communities possess a simpler sulfur cycle than mat communities, reliant on environmental sulfate assimilation, whereas a few orders of the *Alpha*- and *Gamma-proteobacteria* in the salt pans are potentially capable of using sulfate as an electron acceptor for respiration.

### Defense mechanisms against mobile genetic elements are abundant, diverse and novel in Namib Desert saline microbial community metagenomes

To gain further insights into the dynamics of virus-host interactions in the Namib salt pan and halite microbial communities, we assessed the presence of KO terms of the category of Prokaryotic Defense Systems in our metagenomic data. An average of 0.94% of the total KO counts belonged to this category, from which the majority belonged to toxin-antitoxin systems (42.7%), followed by restriction-modification systems (27.8%), CRISPR-Cas systems (23.1%) and DNA phosphorothioation systems (6.4%) (Figure 4A).

**Figure 4.**
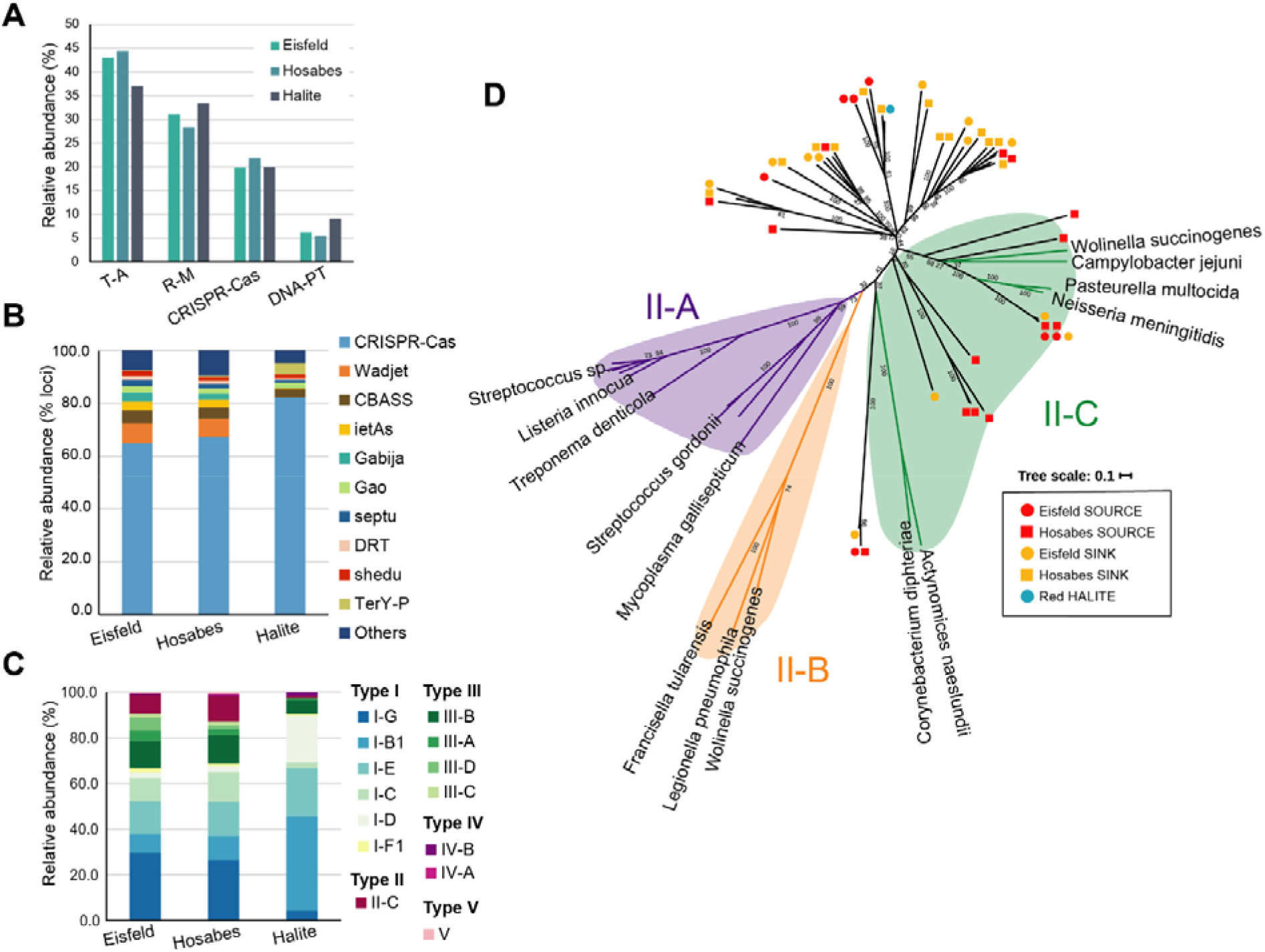
CRISPR-Cas type I and type III systems are widespread in the Namib Desert salt pan microbial communities. **A:** Relative abundance of prokaryotic defense systems KO terms in the Namib Desert stream mat and halite metagenomes. T-A – toxin-antitoxin; R-M – Restriction modification; DNA-PT – DNA phosphorothioation. **B:** Relative abundance of prokaryotic defense systems annotated using the PADLOC. The graph indicates the percentage of total defense loci identified. C: Relative abundance of CRISPR-Cas systems in each of the metagenomic datasets. **D:** Phylogenetic tree of the Cas9 protein sequences present in the Namib salt pan metagenomes. Colored branches indicate a reference protein sequence obtained from the RefSeq protein database, while black branches indicate protein sequences obtained from the metagenomic data. Sequences belonging to the three described Cas9 categories are shaded in purple (II-A), green (II-C) or orange (II-B). The symbols at the end of the branch indicate the location within the salt pan from where the metagenomic sequence was obtained (red: Eisfeld, yellow: Hosabes, blue: halite, circle: source, square: sink).

A detailed search of prokaryotic defense systems was performed with the PADLOC tool (24) to better assess the diversity of defense strategies not covered by the KO database (Supplementary Table 8). The results of both pipelines are complementary since PADLOC focuses on recently discovered defense systems (24) and does not include toxin-antitoxin or restriction modification systems. CRISPR-Cas systems dominated as 66.4% to 87.5% of the detected defense-loci belonged to this category (Figure 4B). The second most-abundant defense system detected was Wadjet, which recognizes extraneous plasmids (25, 26) (Figure 4B). It constituted between 4.8-7.1% of the defense loci in the salt pan metagenomes but was absent from the halite datasets. The cyclic oligonucleotide-based antiphage signaling system (CBASS) was also present in all the metagenomes, ranging from 0.6 to 7.1% of the defense loci (Figure 4B). Other systems constituting on average ≥ 1% of defense loci were ietAS, Gabija, GAO, Septu, Shedu, Zorya and TerY-P (Figure 4B).

CRISPR-Cas adaptive immune systems consist of an array of short sequences (spacers) originating from mobile genetic elements and CRISPR-associated (Cas) proteins required for the acquisition (adaptation) and utilization of spacer sequences and targeting of the invading mobile genetic element (27). Given that the discovery of new CRISPR-Cas systems with unique capabilities is important for the development of new tools with biotechnological application (28), we further investigated the diversity of the salt pan CRISPR-Cas systems (Supplementary Tables 8-9). The majority of the CRISPR-Cas related loci could not be assigned to a functional module and these corresponded to 35.4%-53.4% of all defense loci; additionally, adaptation modules for spacer acquisition represented 17.1%-26.5% of the loci (Figure 4B). The analysis of classified CRISPR-Cas loci in the Namib salt pan and halite metagenomes revealed that type I (which specifically degrade DNA) and type III (which can target both DNA and RNA) CRISPR systems were dominant; representing on average 72.3% and 17.9% of all identified CRISPR systems, respectively (Figure 4C). Type II CRISPR systems (which target DNA) also represented 8.1% of the dataset. Most of the type II CRISPR-marker gene Cas9 sequences belonged to the *Planctomycetota* (36%) and *Verrucomicrobiota* (14.1%) phyla as well as the *Alpha*-(18.4%), *Gamma*-*proteobacteria* (4.4%) and *Acidithiobacillia* (3.2%) pseudodomonadotal classes (Supplementary Table 10). The high representation of cas9 genes from the recently defined sulfur-oxidizing autotrophs *Acidithiobacillia* (29) was remarkable, since this taxon represented from only 0.02% (dark halite) to 0.142% (Hosabes sink) of the total sequences. While the taxonomic distribution of CRISPR-Cas systems is highly patchy - particularly in bacteria -, this result suggests an enrichment of type II CRISPR-Cas systems in the *Acidithiobacillia*. The majority of the Cas9 protein sequences were found to branch deeply within the II-C subtype, with no affiliations to the II-A and II-B subtypes (Figure 4D). These results suggest that a large proportion of the saline sample metagenomic Cas9 sequences may correspond to a novel subtype.

### Viral diversity of the Namib salt pan and halite metagenomes

The VirSorter tool (30) was used to extract putative viral genomic content from the assembled metagenomic data. A total of 3448 contigs were predicted to be of viral origin, of which 857 were over 10 kb in length (Supplementary Table 11). This dataset is subsequently referred to as *mVir*. To compare the similarity of mVir viral populations to viruses in the RefSeq database and previously studied viruses from the Namib (named as NamibVir), a genome-based gene-sharing network was constructed using the vContact 2.0 pipeline which employs a distance-based hierarchical clustering approach to classify viral sequences into clusters that are equal to viral genera (31). The organization of sequences in a network implies a common phylogenetic origin, as occurs for the Caudoviricetes network (Figure 5 inset) (31).

**Figure 5.**
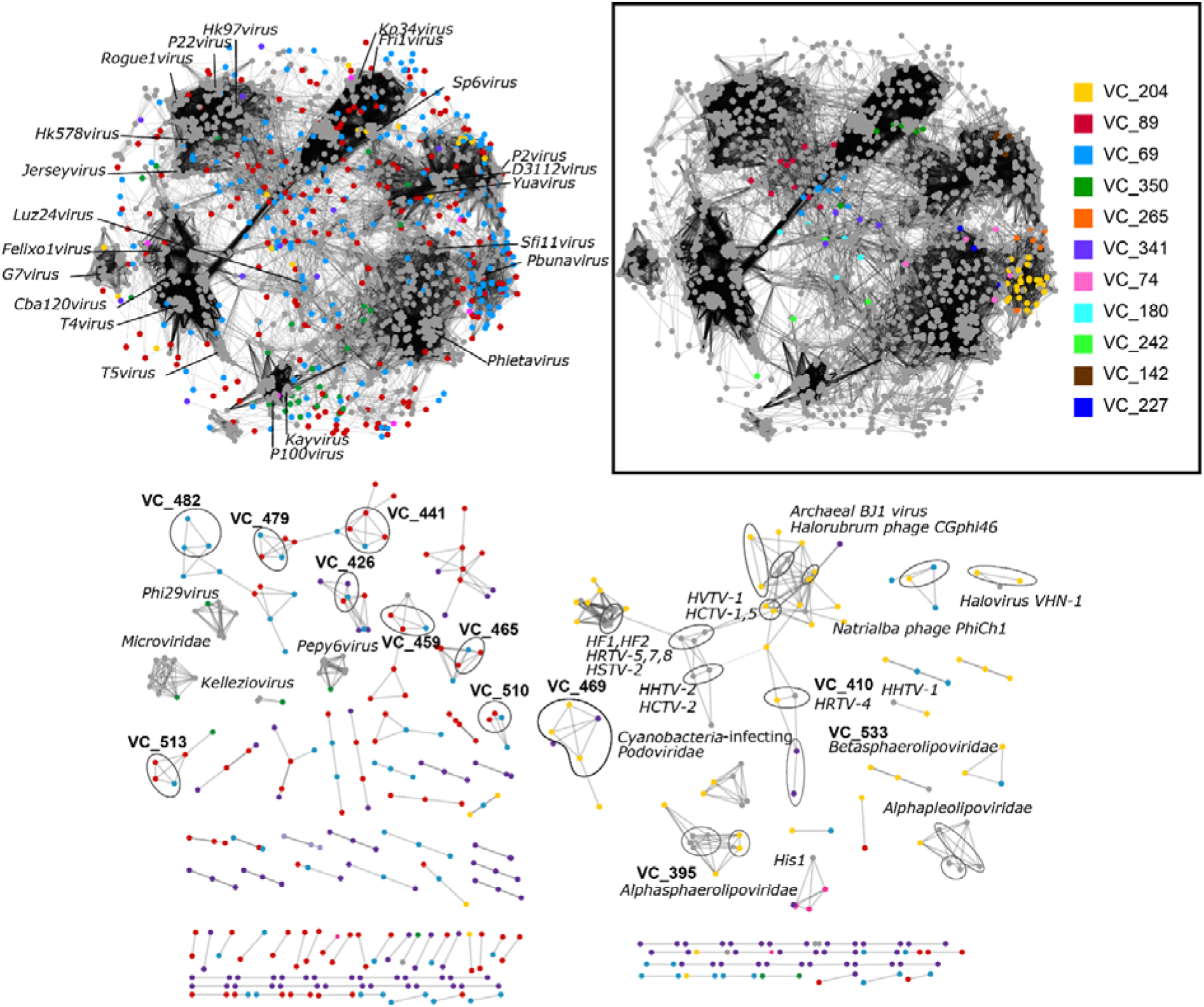
Genome-based network of shared protein content. Each node represents a viral genome and edges represent statistically significant relationships between the protein profiles of those viral genomes. Groups composed exclusively of RefSeq viruses were excluded for clarity. Viral clusters of interest are indicated with the prefix “VC” followed by their corresponding number, and/or circled in black. *Inset:* Network of *Caudoviricetes* indicating the largest viral clusters with mVir sequences.

This analysis resulted in the identification of 145 clusters containing 371 mVir sequences, plus 255 sequences classified as outliers (i.e., with only weak connections to any given cluster and with insufficient information to accurately assign them to genus level) and 197 as singletons (sequences with no similarity to any other, thus not included in the network). The resulting network of clustered viruses is shown in Figure 5. A total of 84% of the mVir clusters (122) were constituted exclusively by mVir sequences, constituting up to 138 putative new viral genera and highlighting the very high genetic diversity of the Namib salt pan viruses (Figure 5 and Supplementary Table 12-14).

Taxonomy assignment of the mVir-containing clusters (*n* ≥3 sequences) revealed that 53 clusters belong to the class Caudoviricetes (Figure 5, inset and Supplementary Table 15), confirming the already observed dominance of this viral order in many Namib Desert niches (13, 32, 33). Only 11 clusters could be classified at family level and 4 at genus level (Supplementary Table 15); three halite virus clusters belonging to the *Halobacteriota-*infecting *Betapleolipovirus* and one to an unclassified halovirus family (Figure 5). Additionally, 23 sequences from the halite mVir metagenomic data formed a cluster together with 17 Halobacteriales*-*infecting viruses from the RefSeq database and 3 NamibVir sequences (Figure 5, *Halovirus* network). Nevertheless, the majority of the mVir sequences could not be clustered at genus level, suggesting they belong to novel archaeoviral taxa infecting Halobacteriales. Other halite mVir sequences were connected to archaeal viruses of the families *Alphasphaerolipoviridae* and *Alphapleolipoviridae*, to the *Caudovirales* main network and to a group of Cyanobacteria*-*infecting *Podoviridae* known to infect freshwater and thermophilic members of this bacterial phylum (Figure 5).

We also compared our mVir dataset to the viral fraction of halite nodules from the hyperarid Atacama Desert (denoted as Hvir; (14)). Only two Hvir sequences clustered together with mVir data, specifically with cluster VC_469 which contained 3 halite mVir and one NamibVir contigs (Supplementary Figure 3 and Supplementary Table 16). Despite the small number of sequences available, the formation of clusters comprised exclusively of metagenomic sequences show that halite rocks harbor novel viral diversity.

### Virus-host interactions reveal novel Planctomycetes-infecting viruses

To investigate virus-host associations, we used an established *in silico* approach based on CRISPR spacer matches between the cellular and viral sequences (34, 35). A total of 1431 CRISPR spacers originating from the salt pan metagenomic data were matched to viral sequences retrieved from the RefSeq, NamibVir and mVir databases (Supplementary Figure 5 and Supplementary Table 17). The majority of the virus-host matches (88.4%) targeted mVir viruses, while 6.6% matches belonged to sequences from one of the NamibVir soil viromes (32) and 4.9 % to RefSeqABV or RefSeq virus databases. Surprisingly, no hits to contigs with taxonomy assignment were found to viral sequences from a previously sequenced metavirome from the same Namib salt pans (13).

A total of 72% of the CRISPR spacer matches arose from the abundant *Pseudomonadota* phylum, of which 43% originated from the *Gammaproteobacteria,* followed by a 7.5% from *Bacteroidetota* and 5.5 % to *Planctomycetota.* Matches to other low abundance taxa were found for the *Cyanobacteria*, Lentisphaerota, *Deltaproteobacteria*, *Gemmatimonadota* and few hits to the halite *Halobacteriota* (Supplementary Figure 5 and Supplementary Table 17).

An alternative approach to establish virus-host linkages is the prediction of proviral sequences within cellular contigs (34). 201 sequences were identified as proviruses of which 119 (59.2%) could be phylogenetically assigned, particularly to *Alphaproteobacteria* (40%), *Planctomycetota* 11% and *Halobacteriota* (10%) contigs (Supplementary Table 18). We note that proviruses were also identified in members of the *Verrucomicrobiota* phylum, where this host linkage was not identified through CRISPR spacer matches. Furthermore, 7 viral clusters (VC_204, VC_69, VC_350, VC_180, VC_469, VC_409 and VC_406), including the largest mVir cluster VC_204, contain prophage sequences, implying a temperate infection mode for these viruses (Table 2).

**Table 2.**
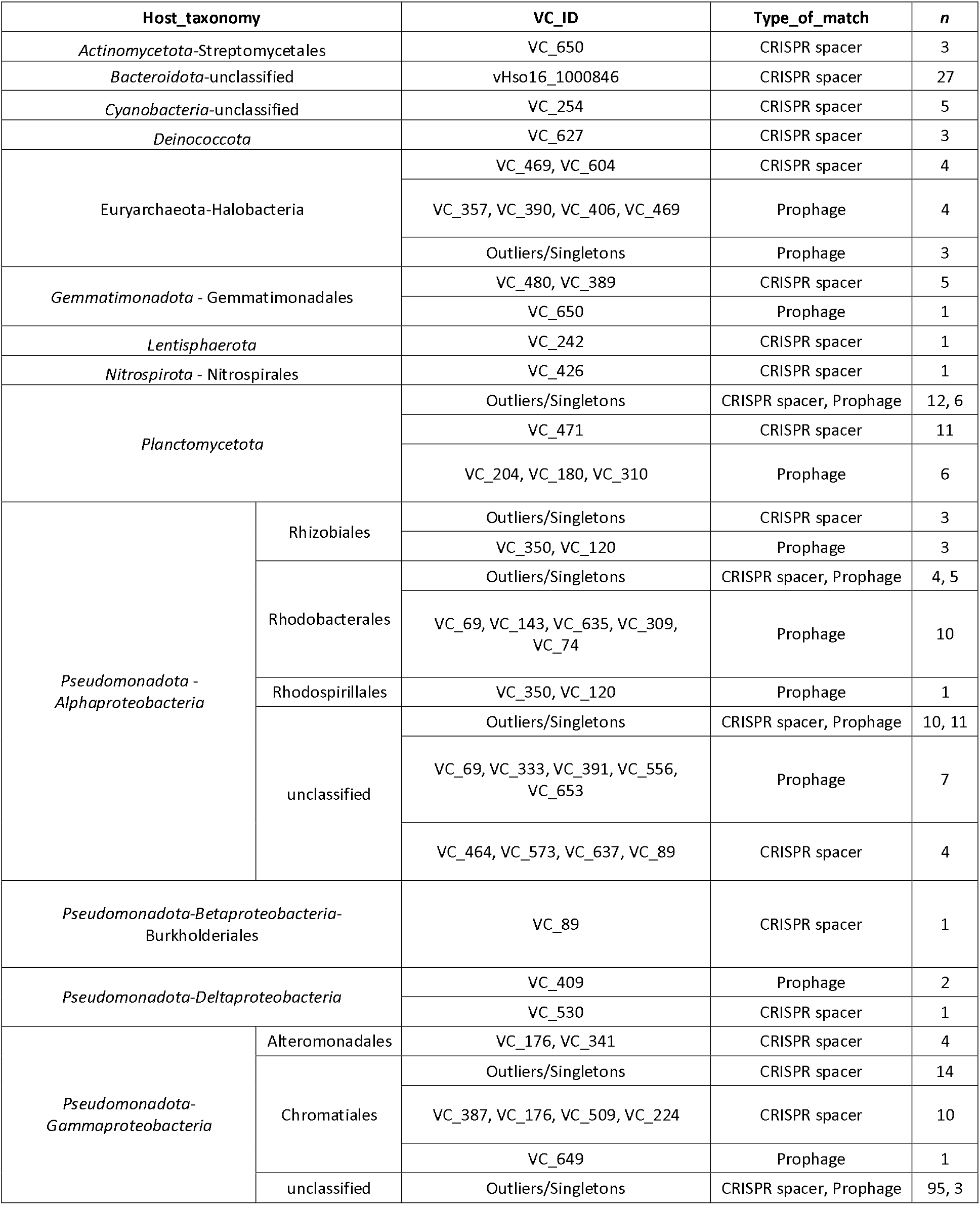
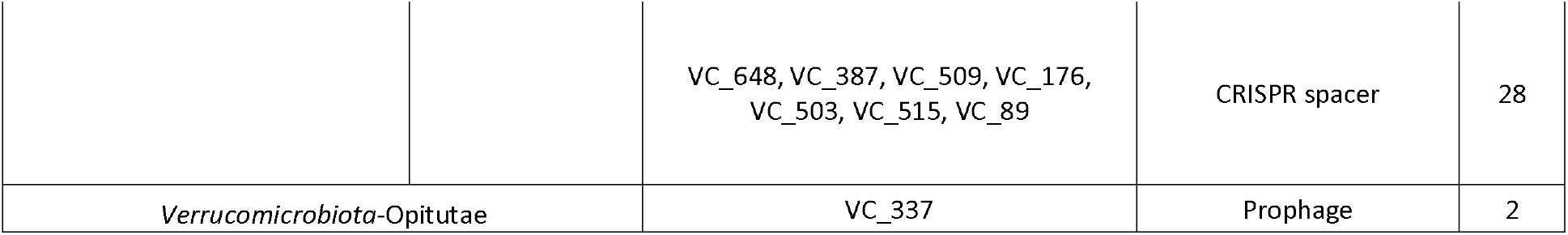
Virus-host connectivity. . Total number of linkages (n) between mVir viral clusters and their predicted hosts.

With the results from CRISPR spacer matches and provirus prediction we were able to identify viral host-interactions for 8 of the largest Caudoviricetes and 4 Euryarchaea-infecting viral clusters. The clusters VC_69 and VC_74 are linked to the alphaproteobacterial Rhodobacterales order, and VC_ 350 to the Rhizobiales and Rhodospirillales ones. VC_204 and VC_180 are connected to the Planctomycetes, and VC_242, to the phylum Lentisphaerae. Most halovirus-host linkages were found through prophage identification, linking clusters VC_357, VC_390, VC_406 and VC_469 to members of the Halobacteriales, with cluster VC_469 specifically linked to the euryarchaeal *Halorubrum* genus (Table 2).

Interestingly, the two most abundant salt pan taxa, i.e., *Alpha*- and *Gamma-proteobacteria*, show different virus-host linkage profiles: the *Gammaproteobacteria* virus-host linkages were mainly identified through CRISPR hits, while those of the *Alphaproteobacteria* were identified through the prediction of prophages. Moreover, viruses linked to the *Gammaproteobacteria* did not belong to the major mVir clusters in the gene-sharing network, while 3 of the 10 largest mVir clusters infected the *Alphaproteobacteria*. Overall, this suggested that stream mat *Gammaproteobacteria* viruses have predominantly a lytic infection mode while the *Alphaproteobacteria* viruses are rather lysogenic.

The low number of CRISPR spacer matches to a previously published Hosabes and Eisfeld saltern metaviromes was unexpected (13). To better explain this finding, an extended network including all the NamibVir sequences regardless of contig length was generated (Supplementary Figure 5). Ninety percent of the metaviromic saltern contigs clustered together in a group of ssDNA viruses of the *Microviridae* family, as reported previously for this dataset (13) (Supplementary Figure 5 and Supplementary Table 19). Given the small size of these viruses (around 5 kb), they were not included in the previous protein-sharing network that only incorporated sequences over 10 kb. Moreover, the *Microviridae* cluster included only one mVir contig. Very few mVir contigs grouped with other metaviromic saltern contigs of putative dsDNA viruses. This demonstrates that the two datasets of viral sequences from the Namib salt pans, the metagenomic mVir and the metaviromic saltern fraction of the NamibVir, represent different populations of viruses with distinct taxonomic affiliations.

### Alphaproteobacterial gene transfer agent-like islands are present in the Namib Desert saline stream mats

From the 43100 mVir ORFs, 44% were functionally annotated; the majority (646/7130) corresponding to proteins involved in DNA metabolism and viral structural proteins (Supplementary Table 20). Transposases and integrases were also especially abundant, accounting for 1.5% and 1.6% of the total ORFs, respectively.

Interestingly, 77 ORFs were annotated as gene transfer agent-like (GTA-like) structural proteins. Gene transfer agents are virus-like particles encoded and produced by their prokaryote hosts and containing random fragments of the host’s genome. Consequently, they are considered to be viable vectors for horizontal gene transfer (36). Given that GTAs are well-documented elements of *Alphaproteobacteria* genomes (36), the presence of GTAs in the mVir data was investigated by performing a protein blast between mVir ORFs and reference GTAs (Supplementary Table 21). A total of 437 mVir contigs matched known *Alphaproteobacteria* GTA proteins. Although 76% of the hits were to unclustered contigs (i.e., contigs unassigned to a viral cluster), 42 clusters matched GTA proteins, including all contigs from clusters VC_69 and VC_89 (Figures 5 and 6). We particularly note that VC_69 harbors 13.8% of the identified prophages that are taxonomically assigned to a viral genus. As GTAs are thought to derive from lysogenic viruses (36), the lysogenic nature of VC_69 supports a link between these viral elements and GTAs.

**Figure 6.**
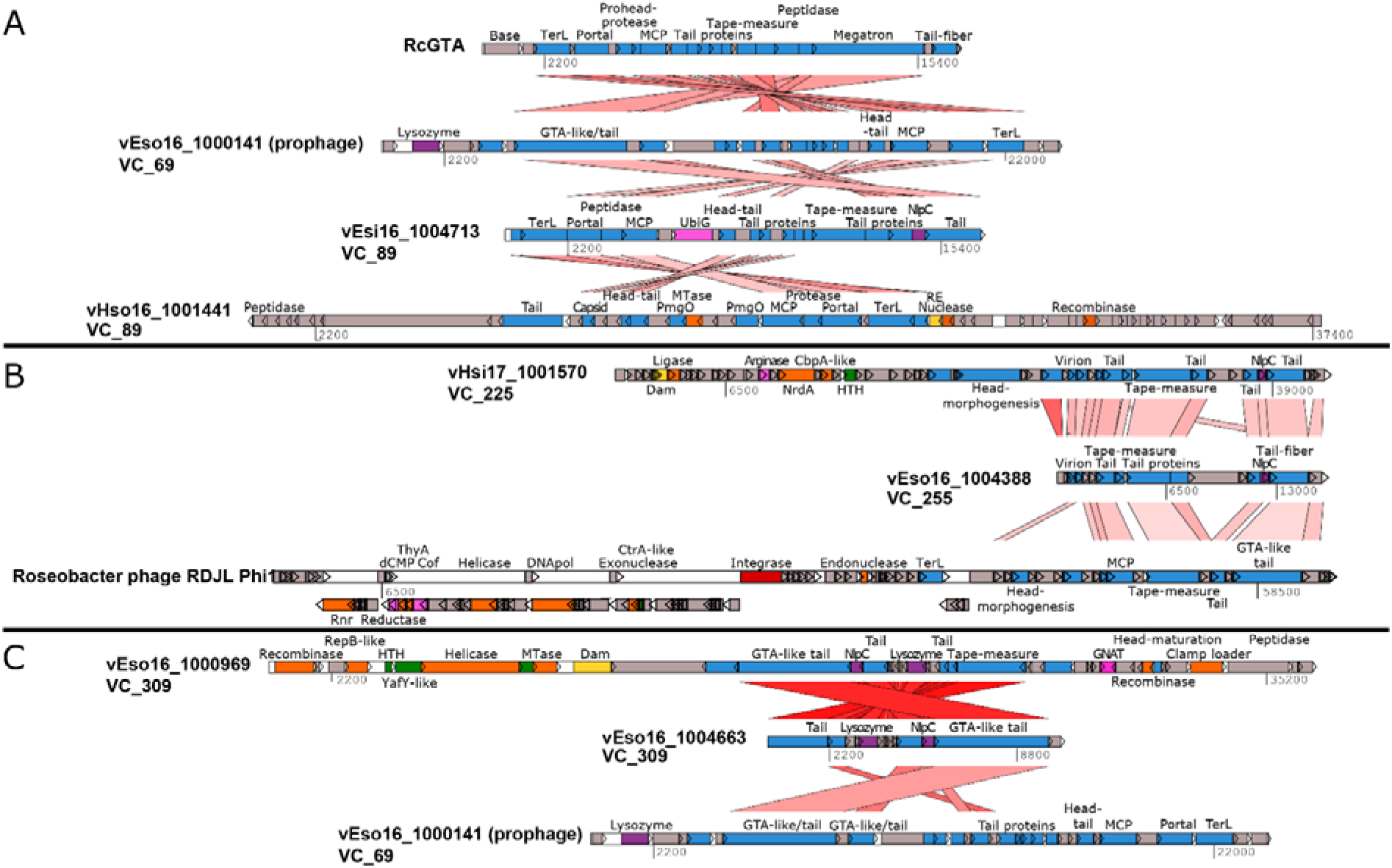
Overview of the genomic organization of GTA-like and virus-like contigs from mVir clusters with GTA-signal. **A**: Comparison of *Rhodobacter capsulatus* GTA (RcGTA) to the largest contig of VC_69 and GTA-like or virus-like contigs from VC_89. **B**: Comparison of VC_255 contigs to *Roseobacter* phage RDJL Phi1 virus. **C**: Genomic organization of VC_309 contigs. The shades of red represent pairwise protein similarity, with stronger red as the most similar. Genes are shown as boxes with an arrow indicating their orientation in the genome and colored according to their assigned functional category: blue – virus structure and assembly, orange – DNA replication, green – transcription, purple – virus exit, yellow – defense mechanisms, red – integration, pink – metabolism.

Protein homology to a GTA ORF is not sufficient to classify a viral sequence as a GTA (36). To distinguish a GTA from a prophage, it should not contain a viral replication module or the small subunit terminase and should have a size of 13-15kb (36, 37). These features were used to screen mVir clusters with a GTA-like signal (Supplementary Table 12; Figure 6 and Supplementary Figure 6). VC_69 exhibited all the necessary GTA-like features (Figure 6A). Moreover, the flanking regions of VC_69 sequences showed a high level of conservation, which would not be expected for “junk” sequences such as defective prophages (Supplementary Figure 7). In the gene-sharing network (Figure 5), VC_69 was connected to the mVir clusters VC_225, VC_309, VC_333 and VC_89, all of which have hits to GTA proteins, and to *Roseobacter phage RDJL1,* a virus phylogenetically related to the *Rhodobacteri capsulatus* GTA (RcGTA) (36). While no replication module was identified in the sequences of cluster VC_69, replication and structural modules were clearly present in VC_225, VC_309 and VC_89 (Figure 6A-C), and the average contig length of these clusters was 22 kb. We note that these clusters contain both GTA-like sequences and sequences that resemble true viruses, contrary to VC_69. Our conclusion is that the latter is better described as a prophage remnant putatively converted into a gene-transfer agent, thus having a potential implication in driving gene exchange in the Namib salt pan *Alphaproteobacteria*.

## Discussion

### Alpha- and Gamma-Proteobacteria dominate the Namib salt pans while Euryarchaea and Salinibacter spp. prevail in the halites

The microbial stream mats from the Namib Desert Hosabes and Eisfeld salt pans showed similar taxonomic and functional profiles, despite being 124 km distant (Figures 1 and 2). This is in agreement with a previous metaviromic analysis reporting that virus communities of both stream mats were also closely related (13). The similarities in the geological and physicochemical composition of the Hosabes and Eisfeld playas, as well as the existence of an underground water system connecting them (6, 7, 13) may explain the taxonomic and functional resemblance of both stream mat microbiomes. Conversely, halite communities were markedly different from the stream mat communities, despite being separated by only a few meters, the disparity being the consequence of the contrasting physicochemical features of each niche, i.e., the mildly saline aquatic mat habitat vs. hypersaline and water-limiting halite; a clear example of environmental filtering (38).

*Pseudomonadota,* in particular *Alpha*- and *Gamma-proteobacteria* were abundant in the salt pan mats (Figure 2B). A dominance of *Pseudomonadota* has been previously reported as a characteristic of saline mat microbiomes (39). The taxonomic diversity profile at phylum level of the stream microbial mats is similar to other saline environments, such as marine water or saline ponds (9, 10, 40–44). However, saline environments can vary widely in their microbial diversities (45). While *Cyanobacteria*, *Pseudomonadota*, *Bacteroidota* and *Chloroflexota* are common members of phototrophic microbial mats (46), the Namib Desert saline spring mats harbor a high percentage of *Planctomycetota*. This phylum has also been identified in hypersaline mats from Shark Bay (Australia), Eleuthera (Bahamas) and Tebenchique lake (Chile) (10, 40, 41), but not in salt pan mats from the Kalahari Desert in southern Africa (47).

Halite microbial communities were markedly different to those inhabiting the surrounding stream mats (Hosabes). Specifically, the triad of microorganisms dominating the halites (the archaeal Halobacteriales order, and the bacterial *Salinibacter* and *Halothece* genera) is almost absent from the mats (Supplementary Tables 3-4). Enrichment of “salt-in” strategists such as the Halobacteriales and *Salinibacter* has been reported in hypersaline environments (9, 11, 18, 42) as this characteristic makes them especially adapted to these habitats. Furthermore, the predominance of the cyanobacterium *Halothece* has also been reported in halite microbial communities from the Atacama Desert (Chile) and the Boneville Salt Flats (USA) (14, 48–52). Overall, the cosmopolitan distribution of these three genera points to highly specialized functional adaptations to saline extremes and possibly also to interactions between them.

### Contrast between the complex biogeochemical cycles of the Namib salt pan mats vs the simple cycles of the oligotrophic halites

*Pseudomonadotas* are possibly key to the functioning of Namib salt pan mats as these were found to have the genetic capacity to perform several steps of the C, N and S biogeochemical cycles. This metabolic diversity has been proposed as a feature that allows them to occupy different trophic niches and survive under fluctuating extreme conditions (39). Although less abundant than the *Pseudomonadotas,* the mat *Planctomycetotas* possess a diverse functional profile (e.g., potential to carry on carbon fixation through the Wood-Ljungdahl pathway, fermentation, nitrification and dissimilatory nitrate reduction) (Figure 3), potentially positioning them as a core taxon for biogeochemical cycling in saline mat communities.

The analysis of the stream mat functional capacity positions the *Cyanobacteria* and the *Alpha*- and *Gamma*- *proteobacteria* as the main primary producers via their photosynthetic capacity, with the additional contribution of *Deltaproteobacteria* and *Planctomycetota* (Figure 3). In contrast, halite carbon fixation was almost an exclusive capacity of the *Cyanobacteria*, particularly to the genus *Halothece*. The latter was not detected in the salt pan mat metagenomes, which strongly suggests that this taxon is highly adapted to hypersaline conditions (Figure 3, (14, 51)). Additionally, the capacity to obtain energy from light (phototrophy) and from CO oxidation (carboxydovory) was widespread in both mat and halite communities, implying that the use of alternative energy sources are key to the hypersaline and oligotrophic desert ecosystem functioning (53).

The limited apparent capacity for nitrogen fixation in the Hosabes and Eisfeld microbial mat assemblages and its absence in the halites suggests these communities may rely on the assimilation of nitrate compounds; particularly in the halites where nitrate assimilation was the main step of the nitrogen cycle for the whole community (Figure 3; (54)). This confirms results obtained from Atacama Desert halites (49) and strongly suggests it is a direct adaptation to the fact that drylands represent the largest terrestrial nitrate pool (54). A probable source of nitrogen to sustain the nitrogen cycle in the Namib playas may be humberstonite (K_3_Na_7_Mg_2_(SO_4_)_6_(NO_3_)_2_*H_2_O), a sulfate-nitrogen mineral that has been only identified in the hyperarid Atacama and Namib Deserts (6).

Similarly, the sulfur cycle of the Namib salt pan mats and halites was potentially dependent on assimilatory sulfate reduction, although capacity for anaerobic sulfate respiration was detected for members of the *Pseudomonadota.* In this regard, the abundant presence of gypsum deposits of the Namib salt pans may represent the source of sulfate for the salt pan microbial mat and halite communities (6).

The main contrast between the stream mat and halite biogeochemical cycles resides in the nitrogen and sulfur cycles, with halite communities having simplified functional capacity relying on assimilation of compounds. These differences could arise from the decrease in diversity associated to the hypersaline conditions of halite minerals. Interestingly, halite minerals from the Atacama Desert have similar taxonomic and functional profiles to the Namib halites. In particular, the absence of nitrogen-fixing *Cyanobacteria* has been reported by several studies (14, 49, 50) and recent work has described simple nitrogen and sulfur cycles limited to the uptake of inorganic nitrogen and sulfur (50). The overall global taxonomic and functional resemblance of halite microbial communities points to the selection of universal specialists adapted to the oligotrophic, hypersaline conditions of halites.

### Putative novel type II CRISPR-Cas systems in the Namib salt pans

The relative abundance of defense systems in the metagenomic data is in accordance with previous reports of the abundance of these systems in bacterial and archaeal genomes, where toxin-antitoxin and restriction modification systems are the most widely distributed and occupy the largest fraction of the genome (55, 56). This applies as well to the proportion of CRISPR-Cas systems, where the abundance of type I (66.7-68.6%) and type III (23.9-18.3%) systems in the stream mat communities follows closely the distribution of CRISPR-Cas in prokaryotic genomes where type III *loci* represent around 25% of all CRISPR-Cas systems and type I CRISPR-Cas systems are the most widespread, with a relative abundance of approximately 60% (57). The high proportion of CRISPR-Cas in halite communities (82.2 %) correlates with the previously estimated abundance of these systems in archaea, where 85.2% of the genomes contain CRISPR-Cas loci and I-B is the most abundant subtype (58).

Although the fraction of type II CRISPR-Cas systems in the salt pan metagenomic data was similar to the previous estimation of type II *loci* abundance in bacterial genomes (57), a phylogenetic analysis of the Cas9 protein, an effector and marker gene of type II systems, reveals novel diversity of sequences branching deeply within Cas9 II-C subtype, putatively constituting a novel subtype of Cas9 proteins (Figure 4D). These new sequences belong to phyla where few type II CRISPR-Cas systems have been previously described, such as the *Planctomycetota* and *Verrucomicrobiota*, underlying the importance of studying uncultured microorganisms of diverse environments. Given the importance of type II CRISPR-Cas systems for genome editing applications (59), these results suggest that the Namib Desert salt pans, as well as other saline systems worldwide, represent an important resource for identification of new CRISPR systems. Indeed, in such environments, where eukaryotes are almost absent, viruses are the main regulators of prokaryote abundances (60).

### Lysogenic viruses infect the main microbial taxa in the Namib salt pan and halite communities

The application of an *in silico* approach to study the Namib salt pan virus population allowed the identification of 138 putative novel viral genera, almost exclusively belonging to the class Caudoviricetes, the largest viral taxon of prokaryotic viruses to date (31) (Supplementary Table 15). Comparisons of the mVir dataset obtained from this study to other viruses from the Namib Desert reveals a niche-dependent viral taxonomic diversity (Figure 5, Supplementary Figure 5), in agreement with the taxonomic differences in the microbial populations inhabiting each type of niche (38).

Novel viruses of the *halobacteriotal* Halobacteriales order were also identified (clusters VC_357, VC_390, VC_406 and VC_460, Figure 5). The addition of mVir data and halite viruses from the Atacama Desert to the known Refseq haloviruses produced a rearrangement in the taxonomic affiliation of some RefSeq viruses, a phenomenon that indicates that haloarchaeal viruses are under-sampled (31) and that further sampling of these viral populations is necessary to better chart these archaeal viruses.

Virus-host linkages were identified for eleven different prokaryotic phyla, especially for the most abundant *Alpha*- and *Gamma-proteobacteria* (Table 2). Interestingly, linkages to *Alphaproteobacteria* were mainly through the identification of proviruses, while those to the *Gammaproteobacteria* were principally through CRISPR spacer hits. This could reflect a divergent infection mode of the viruses infecting each taxon, i.e., lysogenic viruses for *Alphaproteobacteria* and lytic viruses for *Gammaproteobacteria*. The targeting of an integrated element by the CRISPR system would be strongly selected against and could explain the lack of virus-spacer hits in the *Alphaproteobacteria*.

The profuse host associations between the most abundant novel mVir viral genera in the salt pan mats, and the bacterial *Pseudomonadota*, *Planctomycetota* and *Lentisphaerota* phyla, as well as to the halite archaeal Halobacteriales and Haloferacales orders (Table 2), hints at an important role of viruses in nutrient recycling and prokaryotic population control in the Namib salt pan communities through the infection and lysis of the abundant host taxa (61). Additionally, the identification of putative lysogenic viral lineages that include the largest mVir cluster identified in this study (VC_204 and VC_180) and infect members of the *Planctomycetotas* (Supplementary Tables 17-18) suggests that these viruses could impact microbial mat function and structure, since members of this phylum are among the most abundant in the salt pan mats studied and possess unique metabolic capacities within their community (Figures 2B and 3).

One surprising observation of this study is the dissimilar viral taxonomic profiles of the mVir data from this work and a previous metaviromic study of the Hosabes and Eisfeld microbial mat viral populations (13). We conjecture that this could be the result of the different methodological approaches employed to analyze the viral fractions in these communities, where mVir sequences mined from metagenomic data may be enriched in proviruses while extracellular viruses used to produce metaviromes are enriched in lytic virus progeny, as has been suggested by previous work with soil viromes (62, 63). For example, lytic archaeal viruses of the *Salterprovirus* genus (of which virus His1 is the reference strain) (64) were present in the salt pan metavirome but absent in the mVir data, which instead contained several proviruses (Table 2). Additionally, methods associated to metavirome library preparation, specifically the use of multiple displacement amplification (MDA), introduce a strong bias in favor of the amplification of ssDNA, which may explain the overwhelming presence of ssDNA *Microviridae* genomes in the metaviromic salt pan study (65). Taken together, these observations, and the fact that the genetic materials of viruses can be RNA or DNA, argue strongly in favor of using multiple different methods to obtain complementary information to characterize the virus diversity of any community.

### Virus domestication impacts horizontal gene transfer in the Namib salt pan Alphaproteobacteria

It is hypothesized that *in silico* tools used to predict viruses from bulk metagenomic data may include other elements of the mobilome such as plasmids or relic phages (e.g., gene transfer agents [GTAs] and provirus remnants inserted in the microbial genomes) (30, 62, 66). Within this mobilome, GTAs are of special interest. These small virus-like particles are highly abundant in marine environments, where they have been shown to mediate HGT-events at very high frequencies (i.e., 10^-2^ to 10^-4^ (67, 68)). GTAs arise from the incorporation of lysogenic viruses that become inactive and are recruited or “domesticated” by the cell as tools for HGT. As a result, it is difficult to differentiate them from true virus sequences in environmental metagenomes (36).

Surprisingly, the cluster mVir VC_69 was identified as a gene-transfer agent and not as an authentic viral taxon. This cluster had the highest similarity to RcGTA in the mVir data and displayed all features characteristic of GTAs (Figure 6, Supplementary Figure 7), suggesting that the Namib salt pan Rhodobacterales have the capacity to produce GTAs, as other members of the *Alphaproteobacteria* (36). Conversely, clusters VC_89, VC_255 and VC_309 contained both small GTA-like sequences together with *bona fide* viral sequences over 30 kb and with replication modules, suggesting that they correspond to true, active viruses. Although the true impact of GTA-mediated HGT is not known, it is hypothesized to be crucial for cellular adaptation and evolution, driving the diversification and adaptation of the alphaproteobacterial clades containing them to different environments (66). This has important implications in the adaptability of the Namib salt pan Rhodobacterales.

## Conclusions

Deserts are polyextreme environments and their indigenous microbial communities are particularly subjected to water limitation and oligotrophy. Consequently, deserts are among the global ecosystems with the lowest microbial diversities and abundances (69). However, as shown in this study (Figures 2B and 4), desert saltpans and halites still present a high proportion of uncharted microbial – including viral - diversity. Consequently, in the absence of taxonomical and functional assignments for many of the reads and contigs generated, we are almost certainly underestimating the functional capabilities of such communities.

In deserts, playas/salars/salt pans represent niches with constant water availability and are characterized by high salt concentrations. In contrast, halites are salt-saturated rocks with very limited bioavailable water. In this shotgun metagenomic study, we show that the microbial mats and halites of these saline springs constitute a large assemblage of microbial lineages with a vast metabolic genetic versatility, potentially enabling them to cope with the polyextreme environmental conditions. Analyses of the viral fraction of the salt pan microbiome suggests that these habitats are a hub of novel viruses and viral activity and CRISPR-Cas systems. We particularly advocate the use of multiple technical approaches (e.g., metaviromics, metagenomics and metatranscriptomics) to holistically detect the many RNA and DNA viral genomes possibly present. This is particularly relevant since, in (hyper)saline ecosystems, viruses – more than eukaryotes – control the abundances of the bacterial and archaeal populations.

## Materials and Methods

### Sample collection

Sediment and microbial mat samples were collected under sterile conditions in Whirlpack® bags from two salt pans in the Namib Desert in April 2016 and 2017: Hosabes (S 23°30’425’’, E 15°04’309’’) and Eisfeld (S 22°29’002’’, E 14°34’363’’) salt pans (Figure 1). For each sampling campaign, samples were collected at the ‘source’ and ‘sink’ of each salt pan. Temperature and conductivity measurements were taken on site at the time of sampling. Both years the mat were covered by ∼ 5 cm of stream water and we collected the first 5-8 cm of mat which included sediments. The sampled were collected very close (∼ 20 cm) to each other both years. Samples from two halites close to the stream of the Hosabes salt pan were also collected in 2017: a red (S 23°30’25.1’’, E 15°04’17.1’’; Figure 1D) and a black halite (S 23°30’25.1’’, E 15°04’17.5’’; Figure 1E). The samples were stored at room temperature prior to their transport the CMEG laboratory, where they were stored at −20°C until metagenomic DNA (mDNA) extraction.

### DNA extraction and sequencing

Samples were thawed on ice and 6-10 aliquots of approximately 0.25 g each were subjected to DNA extraction using the PowerLyzer® PowerSoil® DNA Isolation kit (QIAGEN) following the manufactureŕs instructions. Around five extractions were performed per sample. Prior to mDNA extraction, the halite samples were pulverized with a sterile mortar and dissolved in a sterile 20% NaCl solution and the biomass was recovered by centrifugation at 10 000 x g for 15 min, 4°C. Aliquots of approximately 0.2 g of biomass pellets were further used for mDNA extraction also with the PowerLyzer® PowerSoil® DNA Isolation kit (QIAGEN).

DNA samples from the same location were pooled, concentrated using ethanol precipitation (70), and further purified using the DNeasy PowerClean CleanUp kit (QIAGEN). Consequently, ten composite mDNA preparations (from 8 salt pan and 2 halites, respectively) underwent library construction and sequencing by Admera Health, LCC (NJ, USA). The PCR-free KAPA Hyper prep kit was used for library preparation and paired-end reads of 150 bp were sequenced in a single lane using the Illumina HiSeq X platform.

### Quality control, filtering and assembly of the sequenced data

Between 53 (Eisfeld sink 2017) and 132 (Hosabes sink 2016) million paired-end reads were obtained for each sample (Supplementary Table 1). The quality of the reads was checked using FastQC (71). The BBTools package (<u>BBMap – Bushnell B.-sourceforge.net/projects/bbmap</u>) was used for read filtering: adapter trimming was done with BBDuk using the recommended settings (ktrim=r k=23 mink=11 hdist=1 tpe tbo t=10) and the read quality trimming was done at the Q10 quality level, with a minimum average quality of Q15 and a minimum length of 100 bp. The dedupe script (BBTools package) was used with the recommended settings (ac=f) to remove exact duplicates. The post-QC reads were assembled with SPAdes v3.9.0 (72) using the ‘meta’ flag. Only contigs over 500 bp were retained for further analysis. Assembly statistics were calculated with the Stats script implemented in the BBTools package and BBMap was used for the calculation of the sequencing depth per contig. Reads were mapped to the assembly using Bowtie2 (73). The quality filtering and assembly results are shown in Supplementary Table 1.

The 10 resulting contig files were uploaded to the IMG/M system (74) for functional and phylogenetic annotation and are available under the Study ID Gs0133438, Analysis Projects Ga0248485, Ga0248504, Ga0254891, Ga0255014, Ga0255825, Ga0256419, Ga0256679 and Ga0256680.

### Shotgun metagenome phylogenetic and functional analysis

To gain insight into the genetic potential of the Namib salt pan and halite microbial communities, two approaches were combined. KOs belonging to the most abundant taxa (>5 %) were screened for the presence of complete KEGG modules (see Materials and Methods and Supplementary Table 6). Secondly, pathway hallmark genes were used as markers to assess the functional potential of the community, as well as the taxa possessing such genetic potential.

The taxonomic assignment of the genes of each dataset was retrieved from the IMG/M annotation pipeline (74). To obtain the taxonomic profile of each metagenome (Figure 2B), the reads from each metagenome were mapped to the predicted genes and the relative abundance of each gene expressed as transcripts per million (TPM) was calculated as described previously (75). For comparisons of the taxonomic composition of microbial communities (Figure 2A and Table 1) a raw count matrix of taxa at family level was created by combining the gene counts for each family in each metagenome. The resulting matrix was subjected to differential statistical analysis with the R package *DESeq2* (76) using the Walden test with a *Padj ≤* 0.05. Only taxa with a log2FoldChange (log2FC) ≥ 1 were further considered (Table 1).

For functional analysis, the assignment of KO terms to the predicted open reading frames (ORFs) was retrieved from the IMG/M pipeline (74). Phyla with more than 5% of abundance were selected for taxon-based functional analysis. Consequently, only genes assigned to the phyla *Pseudomonadota*, *Bacteroidota*, *Cyanobacteria*, *Planctomycetota* and *Halobacteriota* were retrieved, and the KO terms associated to each taxonomic group submitted to KEGG Mapper (77) for the reconstruction of complete KEGG functional modules. A matrix of KO counts was constructed relating the KEGG modules to the taxa in each metagenome and normalized by the total KO terms of each dataset (Supplementary Table 6). Additionally, metabolic processes were investigated using specific KO terms as markers, and their relative abundances were calculated as described above (Supplementary Table 7).

The identification and annotation of prokaryotic defense systems was done using the tool PADLOC (padlocdb v1.4.0) (24) and the results of the analysis can be found in Supplementary Table 8.

### CRISPR-Cas classification

PSI-BLAST (78) to the Conserved Domains database (CDD) (79) (downloaded on March 2017) with an E value ≤ 10^-6^ was used to identify Cas genes present in the salt pan metagenomic data (57). Multigenic *cas* loci were defined as two or more *cas* genes within 5 genes up- or down-stream of each other. Clusters were classified using weighted consensus of all members, assigning a value to each depending on their subtype specificity following the criteria established previously (57). For type II loci subtype classification, protein sequences of the hallmark Cas9 gene larger than 500 amino acids were extracted and aligned together with reference Cas9 sequences from the Swissprot (2019) and CDD databases using MAFFT (80) with default parameters. Duplicates were removed by CD-HIT (81) and the alignment was refined using MaxAlign (82). A phylogenetic Neighbor-Joining tree was built using the WAG substitution model and 500 bootstrap resamplings for all gap-free sites.

### Identification of viral genomic from the Namib Desert metagenomic datasets

The metagenomic contig datasets were each processed with VirSorter v1.0.3 (February 2018) (30) through the Cyverse Discovery Environment using both the RefSeqABVir and the Virome databases, and as recommended only categories 1, 2, 4 and 5 were kept for further analysis. A total of 3485 predicted viral sequences were obtained from the complete metagenomic data after removal of duplicated entries. Among these, 201 sequences were identified as prophages of which 44 (i.e., 20%) were circular (Supplementary Table 11). Manual curation of the sequences further revealed the presence of 37 small contigs (shorter than 5 kb in length) and of high coverage only composed of genes encoding for integrases, transposases and recombinases. These most probably represent repetitive regions and were not considered for further analysis. The resulting viral database is named as mVir.

### Construction and curation of a Namib Desert virome database

The Namib Desert viruses database (namely, NamibVir) was constructed by retrieving viral sequences from all the Namib Desert metaviromic studies available (i.e., from hypolith, salt pan and soil samples; (13, 32, 33). To remove contaminant microbial sequences, the metaviromic contigs were processed with VirSorter using the virome decontamination mode (30). Ultimately, the NamibVir database contained 57740 sequences of which only 101 had a size of 10 kb or longer.

### Identification of virus-host pairs using CRISPR spacer matches

Taxonomic classification of the CRISPR *loci*-containing scaffolds was obtained using the Contig Annotation Tool (83). Taxonomic classification of the viral sequences was inferred based on the consensus of BLASTP hits (E value ≤ 10^-5^) to the RefSeq virus proteins (84) (downloaded on June, 2018), using MEGAN4 (85) to assign contigs to taxa.

A total of 4821 CRISPR *loci,* containing 38126 spacer sequences, were retrieved from the IMG/M annotation (74) of the salt pan and halite metagenomic datasets. These spacers were used to create a spacer database. To identify virus-host pairs, the spacer database was compared to four different virus databases: RefSeq viral genomes (84), RefSeqABV (30), NamibVir and the predicted metagenomic viral sequences (mVir database) as described previously (35). Briefly: spacers were aligned to viral sequences using blastn (blastn-short task, E-value of 10^-10^, percent identity of 95% and max_target_sequences = 1). This resulted in 1542 virus-spacer linkages of which only the 1145 matches with contig taxonomy assignment were used to construct a map of virus-host interactions that was visualized using Cytoscape (https://cytoscape.org/ v 3.7.0) (Supplementary Table 17).

### Construction of a genome-based viral network

Proteins in the 857 mVir genomes over 10 kb were clustered with proteins from 3747 viral genomes in RefSeq (June 2018) and the 101 genomes (>10 kb) from NamibVir database using an all-versus-all BLASTP (E-value 0.00001) followed by the aggrupation into protein clusters as previously described (86). A similarity score was calculated using vContact v2.0 (31) and the resulting network was visualized with Cytoscape (“https://cytoscape.org/” v 3.7.0) using an edge-weighted spring model. Taxonomy assignment of the mVir viral clusters was done following three criteria: if the cluster included one or more RefSeq viruses their taxonomic affiliation from the NCBI taxonomy was assigned to the viral cluster at the lowest taxonomy rank in common (using a 75% cut-off value). If the cluster consisted exclusively of mVir and NamibVir sequences, the lowest taxonomy rank in common to all sequences (using a 70% cut-off value) obtained from the consensus of BLASTP hits as described above was selected as putative taxonomy of the cluster. Finally, network topology was also used to assign taxonomy. If the sequences belonged to a network of sequences containing RefSeq viruses with taxonomic consensus at order level they were automatically assigned to that order.

### Annotation of viral sequences

The mVir genes were annotated by using a combination of the functional annotation retrieved from the IMG/M pipeline with the result of matching the viral ORFs to the Prokaryotic Virus Orthologous Groups (pVOG) database (87) using hmmsearch (88) with an E-value threshold of 10^-5^.

### BLASTP vs GTAs

BLASTP similarity searches were carried out on all mVir ORFs using the genes of *Rhodobacter capsulatus* (GenBank: AF181080.3) and *Bartonella australis* GTAs (89), retaining matches with E-values ≤ 10^-5^. Results are reported in Supplementary Table 21.

### Accession numbers

Metagenomic data generated in this work can be accessed through the IMG/M database (https://www.img.jgi.doe.gov) under GOLD Sequencing Project ID: Gp0293142 and IMG Genome IDs: 3300023218, 3300023197, 3300022725, 3300023214, 3300023202, 3300022723, 3300022777, 3300022719, 3300022719 and 3300022719. The unassembled reads from all datasets analyzed are available in the SRA database under BioProject PRJNA943124, accession numbers: SRR23862440, SRR23862446, SRR23862438, SRR23862439, SRR23862444, SRR23862445, SRR23862443, SRR23862437, SRR23862441, SRR23862442.

## Acknowledgements

The authors acknowledge funding support from the University of Pretoria and the National Research Foundation (NRF grant 113308). The authors declare that no conflicts of interest exist.

## Author contributions

LMA designed the study, performed the experimental work and bioinformatic analysis of the data and drafted the manuscript. SV was involved in the taxonomic analysis of the contigs and CLS carried out the classification of CRISPR-Cas systems. JBR participated in sample collection and critical revisions of the manuscript. GMK provided logistical support in the Namib Desert. DAC provided funding and analysis tools, and assisted with manuscript revisions. All authors have read and approved the final manuscript.

## References

1. Bryant DA, Frigaard, Niels-Ulrik. 1996. Validated linear mixture modelling of Landsat TM data for mapping evaporite minerals on a playa surface: methods and applications. 2. International Journal of Remote Sensing 17:315–330.

2. Waiser MJ, Robarts RD. 2009. Saline Inland Waters, p. 634–644. In Likens, GE (ed.), Encyclopedia of Inland Waters. Academic Press, Oxford.

3. Eckardt FD, Drake N. 2011. Introducing the Namib Desert Playas, p. 19–25. *In* Öztürk, M, Böer, B, Barth, H-J, Clüsener-Godt, M, Khan, MA, Breckle, S-W (eds.), Sabkha Ecosystems: Volume III: Africa and Southern Europe. Springer Netherlands, Dordrecht.

4. Ward JD, Seely MK, Lancaster N. 1983. On the antiquity of the Namib. South African Journal of Science 79:9.

5. Day J, Seely M. 2004. Physical and chemical conditions in an hypersaline spring in the Namib Desert. Hydrobiologia https://doi.org/10.1007/BF00015477.

6. Eckardt FD, Drake N, Goudie AS, White K, Viles H. 2001. The role of playas in pedogenic gypsum crust formation in the Central Namib Desert: a theoretical model. Earth Surface Processes and Landforms 26:1177–1193.

7. Day JA. 1993. The major ion chemistry of some southern African saline systems, p. 37–59. *In* Hurlbert, SH (ed.), Saline Lakes V. Springer Netherlands.

8. Fourçans A, Solé A, Diestra E, Ranchou-Peyruse A, Esteve I, Caumette P, Duran R. 2006. Vertical migration of phototrophic bacterial populations in a hypersaline microbial mat from Salins-de-Giraud (Camargue, France). FEMS Microbiology Ecology 57:367–377.

9. Benlloch S, López-López A, Casamayor EO, Øvreås L, Goddard V, Daae FL, Smerdon G, Massana R, Joint I, Thingstad F, Pedrós-Alió C, Rodríguez-Valera F. 2002. Prokaryotic genetic diversity throughout the salinity gradient of a coastal solar saltern. Environ Microbiol 4:349–360.

10. Fernandez AB, Rasuk MC, Visscher PT, Contreras M, Novoa F, Poire DG, Patterson MM, Ventosa A, Farias ME. 2016. Microbial Diversity in Sediment Ecosystems (Evaporites Domes, Microbial Mats, and Crusts) of Hypersaline Laguna Tebenquiche, Salar de Atacama, Chile. Front Microbiol 7:1284.

11. Vera-Gargallo B, Chowdhury TR, Brown J, Fansler SJ, Durán-Viseras A, Sánchez-Porro C, Bailey VL, Jansson JK, Ventosa A. 2019. Spatial distribution of prokaryotic communities in hypersaline soils. Sci Rep 9:1769.

12. Dupraz C, Visscher PT. 2005. Microbial lithification in marine stromatolites and hypersaline mats. Trends Microbiol 13:429–438.

13. Adriaenssens EM, van Zyl LJ, Cowan DA, Trindade MI. 2016. Metaviromics of Namib Desert Salt Pans: A Novel Lineage of Haloarchaeal Salterproviruses and a Rich Source of ssDNA Viruses. Viruses 8.

14. Crits-Christoph A, Gelsinger DR, Ma B, Wierzchos J, Ravel J, Davila A, Casero MC, DiRuggiero J. 2016. Functional interactions of archaea, bacteria and viruses in a hypersaline endolithic community. Environ Microbiol 18:2064–2077.

15. Santos F, Yarza P, Parro V, Meseguer I, Rosselló-Móra R, Antón J. 2012. Culture-independent approaches for studying viruses from hypersaline environments. Appl Environ Microbiol 78:1635– 1643.

16. Sime-Ngando T. 2014. Environmental bacteriophages: viruses of microbes in aquatic ecosystems. Front Microbiol 5:355.

17. Brum JR, Sullivan MB. 2015. Rising to the challenge: accelerated pace of discovery transforms marine virology. Nat Rev Microbiol 13:147–159.

18. Oren A. 2008. Microbial life at high salt concentrations: phylogenetic and metabolic diversity. Saline Syst 4:2.

19. Qian Z, Tianwei H, Mackey HR, van Loosdrecht MCM, Guanghao C. 2019. Recent advances in dissimilatory sulfate reduction: From metabolic study to application. Water Research 150:162–181.

20. Cronan JE, Laporte D. 2005. Tricarboxylic Acid Cycle and Glyoxylate Bypass. EcoSal Plus 1.

21. Peyraud R, Kiefer P, Christen P, Massou S, Portais J-C, Vorholt JA. 2009. Demonstration of the ethylmalonyl-CoA pathway by using 13C metabolomics. PNAS 106:4846–4851.

22. Borjian F, Han J, Hou J, Xiang H, Berg IA. 2016. The methylaspartate cycle in haloarchaea and its possible role in carbon metabolism. ISME J 10:546–557.

23. Petushkova E, Mayorova E, Tsygankov A. 2021. TCA Cycle Replenishing Pathways in Photosynthetic Purple Non-Sulfur Bacteria Growing with Acetate. Life (Basel) 11:711.

24. Payne LJ, Todeschini TC, Wu Y, Perry BJ, Ronson CW, Fineran PC, Nobrega FL, Jackson SA. 2021. Identification and classification of antiviral defence systems in bacteria and archaea with PADLOC reveals new system types. Nucleic Acids Research 49:10868–10878.

25. Doron S, Melamed S, Ofir G, Leavitt A, Lopatina A, Keren M, Amitai G, Sorek R. 2018. Systematic discovery of antiphage defense systems in the microbial pangenome. Science 359:eaar4120.

26. Deep A, Gu Y, Gao Y-Q, Ego KM, Herzik MA, Zhou H, Corbett KD. 2022. The SMC-family Wadjet complex protects bacteria from plasmid transformation by recognition and cleavage of closed-circular DNA. Molecular Cell 82:4145–4159.e7.

27. Marraffini LA. 2015. CRISPR-Cas immunity in prokaryotes. Nature 526:55–61.

28. Pickar-Oliver A, Gersbach CA. 2019. The next generation of CRISPR–Cas technologies and applications. 8. Nat Rev Mol Cell Biol 20:490–507.

29. Hudson CM, Williams KP, Kelly DP. 2014. Definitive assignment by multigenome analysis of the gammaproteobacterial genus Thermithiobacillus to the class Acidithiobacillia. Pol J Microbiol 63:245– 247.

30. Roux S, Enault F, Hurwitz BL, Sullivan MB. 2015. VirSorter: mining viral signal from microbial genomic data. PeerJ 3:e985.

31. Bin Jang H, Bolduc B, Zablocki O, Kuhn JH, Roux S, Adriaenssens EM, Brister JR, Kropinski AM, Krupovic M, Lavigne R, Turner D, Sullivan MB. 2019. Taxonomic assignment of uncultivated prokaryotic virus genomes is enabled by gene-sharing networks. Nat Biotechnol 37:632–639.

32. Hesse U, van Heusden P, Kirby BM, Olonade I, van Zyl LJ, Trindade M. 2017. Virome Assembly and Annotation: A Surprise in the Namib Desert. Front Microbiol 8:13.

33. Zablocki O, Adriaenssens EM, Frossard A, Seely M, Ramond J-B, Cowan D. 2017. Metaviromes of Extracellular Soil Viruses along a Namib Desert Aridity Gradient. Genome Announc 5.

34. Edwards RA, McNair K, Faust K, Raes J, Dutilh BE. 2016. Computational approaches to predict bacteriophage-host relationships. FEMS Microbiol Rev 40:258–272.

35. Paez-Espino D, Eloe-Fadrosh EA, Pavlopoulos GA, Thomas AD, Huntemann M, Mikhailova N, Rubin E, Ivanova NN, Kyrpides NC. 2016. Uncovering Earth’s virome. Nature 536:425–430.

36. Lang AS, Westbye AB, Beatty JT. 2017. The Distribution, Evolution, and Roles of Gene Transfer Agents in Prokaryotic Genetic Exchange. Annu Rev Virol 4:87–104.

37. Paul JH. 2008. Prophages in marine bacteria: dangerous molecular time bombs or the key to survival in the seas? ISME J 2:579–589.

38. Johnson RM, Ramond J-B, Gunnigle E, Seely M, Cowan DA. 2017. Namib Desert edaphic bacterial, fungal and archaeal communities assemble through deterministic processes but are influenced by different abiotic parameters. Extremophiles 21:381–392.

39. Bolhuis H, Cretoiu MS, Stal LJ. 2014. Molecular ecology of microbial mats. FEMS Microbiol Ecol 90:335–350.

40. Allen MA, Goh F, Burns BP, Neilan BA. 2009. Bacterial, archaeal and eukaryotic diversity of smooth and pustular microbial mat communities in the hypersaline lagoon of Shark Bay. Geobiology 7:82–96.

41. Baumgartner LK, Dupraz C, Buckley DH, Spear JR, Pace NR, Visscher PT. 2009. Microbial species richness and metabolic activities in hypersaline microbial mats: insight into biosignature formation through lithification. Astrobiology 9:861–874.

42. Kimbrel JA, Ballor N, Wu Y-W, David MM, Hazen TC, Simmons BA, Singer SW, Jansson JK. 2018. Microbial Community Structure and Functional Potential Along a Hypersaline Gradient. Front Microbiol 9:1492.

43. Sunagawa S, Coelho LP, Chaffron S, Kultima JR, Labadie K, Salazar G, Djahanschiri B, Zeller G, Mende DR, Alberti A, Cornejo-Castillo FM, Costea PI, Cruaud C, d’Ovidio F, Engelen S, Ferrera I, Gasol JM, Guidi L, Hildebrand F, Kokoszka F, Lepoivre C, Lima-Mendez G, Poulain J, Poulos BT, Royo-Llonch M, Sarmento H, Vieira-Silva S, Dimier C, Picheral M, Searson S, Kandels-Lewis S, Tara Oceans coordinators, Bowler C, de Vargas C, Gorsky G, Grimsley N, Hingamp P, Iudicone D, Jaillon O, Not F, Ogata H, Pesant S, Speich S, Stemmann L, Sullivan MB, Weissenbach J, Wincker P, Karsenti E, Raes J, Acinas SG, Bork P. 2015. Ocean plankton. Structure and function of the global ocean microbiome. Science 348:1261359.

44. Zhang W, Ding W, Li Y-X, Tam C, Bougouffa S, Wang R, Pei B, Chiang H, Leung P, Lu Y, Sun J, Fu H, Bajic VB, Liu H, Webster NS, Qian P-Y. 2019. Marine biofilms constitute a bank of hidden microbial diversity and functional potential. Nat Commun 10:517.

45. Zhang K, Shi Y, Cui X, Yue P, Li K, Liu X, Tripathi BM, Chu H. 2019. Salinity Is a Key Determinant for Soil Microbial Communities in a Desert Ecosystem. mSystems 4:e00225–18.

46. Prieto-Barajas CM, Valencia-Cantero E, Santoyo G. 2018. Microbial mat ecosystems: Structure types, functional diversity, and biotechnological application. Electronic Journal of Biotechnology 31:48–56.

47. Genderjahn S, Alawi M, Mangelsdorf K, Horn F, Wagner D. 2018. Desiccation- and Saline-Tolerant Bacteria and Archaea in Kalahari Pan Sediments. Front Microbiol 9:2082.

48. Davila AF, Hawes I, Araya JG, Gelsinger DR, DiRuggiero J, Ascaso C, Osano A, Wierzchos J. 2015. In situ metabolism in halite endolithic microbial communities of the hyperarid Atacama Desert. Front Microbiol 6:1035.

49. Finstad KM, Probst AJ, Thomas BC, Andersen GL, Demergasso C, Echeverría A, Amundson RG, Banfield JF. 2017. Microbial Community Structure and the Persistence of Cyanobacterial Populations in Salt Crusts of the Hyperarid Atacama Desert from Genome-Resolved Metagenomics. Front Microbiol 8:1435.

50. Gómez-Silva B, Vilo-Muñoz C, Galetović A, Dong Q, Castelán-Sánchez HG, Pérez-Llano Y, Sánchez-Carbente MDR, Dávila-Ramos S, Cortés-López NG, Martínez-Ávila L, Dobson ADW, Batista-García RA. 2019. Metagenomics of Atacama Lithobiontic Extremophile Life Unveils Highlights on Fungal Communities, Biogeochemical Cycles and Carbohydrate-Active Enzymes. Microorganisms 7.

51. de Los Ríos A, Valea S, Ascaso C, Davila A, Kastovsky J, McKay CP, Gómez-Silva B, Wierzchos J. 2010. Comparative analysis of the microbial communities inhabiting halite evaporites of the Atacama Desert. Int Microbiol 13:79–89.

52. McGonigle JM, Bernau JA, Bowen BB, Brazelton WJ. 2019. Robust Archaeal and Bacterial Communities Inhabit Shallow Subsurface Sediments of the Bonneville Salt Flats. mSphere 4.

53. Cordero PRF, Bayly K, Man Leung P, Huang C, Islam ZF, Schittenhelm RB, King GM, Greening C. 2019. Atmospheric carbon monoxide oxidation is a widespread mechanism supporting microbial survival. 11. The ISME Journal 13:2868–2881.

54. Ramond J-B, Jordaan K, Díez B, Heinzelmann SM, Cowan DA. 2022. Microbial Biogeochemical Cycling of Nitrogen in Arid Ecosystems. Microbiol Mol Biol Rev 86:e0010921.

55. Koonin EV, Makarova KS, Zhang F. 2017. Diversity, classification and evolution of CRISPR-Cas systems. Curr Opin Microbiol 37:67–78.

56. Puigbò P, Makarova KS, Kristensen DM, Wolf YI, Koonin EV. 2017. Reconstruction of the evolution of microbial defense systems. BMC Evol Biol 17:94.

57. Makarova KS, Koonin EV. 2015. Annotation and Classification of CRISPR-Cas Systems. Methods Mol Biol 1311:47–75.

58. Makarova KS, Wolf YI, Iranzo J, Shmakov SA, Alkhnbashi OS, Brouns SJJ, Charpentier E, Cheng D, Haft DH, Horvath P, Moineau S, Mojica FJM, Scott D, Shah SA, Siksnys V, Terns MP, Venclovas Č, White MF, Yakunin AF, Yan W, Zhang F, Garrett RA, Backofen R, van der Oost J, Barrangou R, Koonin EV. 2020. Evolutionary classification of CRISPR–Cas systems: a burst of class 2 and derived variants. 2. Nat Rev Microbiol 18:67–83.

59. Lau C-H. 2018. Applications of CRISPR-Cas in Bioengineering, Biotechnology, and Translational Research. The CRISPR Journal 1:379–404.

60. Ramos-Barbero MD, Martínez JM, Almansa C, Rodríguez N, Villamor J, Gomariz M, Escudero C, Rubin S dC, Antón J, Martínez-García M, Amils R. 2019. Prokaryotic and viral community structure in the singular chaotropic salt lake Salar de Uyuni. Environ Microbiol 21:2029–2042.

61. van Zyl LJ, Alvarez LM, Trindade M. 2022. Journey of a Thousand Miles: The Evolution of Our Understanding of Viruses in Hot Deserts, p. 133–160. In Ramond, J-B, Cowan, DA (eds.), Microbiology of Hot Deserts. Springer International Publishing, Cham.

62. Emerson JB, Roux S, Brum JR, Bolduc B, Woodcroft BJ, Jang HB, Singleton CM, Solden LM, Naas AE, Boyd JA, Hodgkins SB, Wilson RM, Trubl G, Li C, Frolking S, Pope PB, Wrighton KC, Crill PM, Chanton JP, Saleska SR, Tyson GW, Rich VI, Sullivan MB. 2018. Host-linked soil viral ecology along a permafrost thaw gradient. Nat Microbiol 3:870–880.

63. Trubl G, Jang HB, Roux S, Emerson JB, Solonenko N, Vik DR, Solden L, Ellenbogen J, Runyon AT, Bolduc B, Woodcroft BJ, Saleska SR, Tyson GW, Wrighton KC, Sullivan MB, Rich VI. 2018. Soil Viruses Are Underexplored Players in Ecosystem Carbon Processing. mSystems 3.

64. Bath C, Cukalac T, Porter K, Dyall-Smith ML. 2006. His1 and His2 are distantly related, spindle-shaped haloviruses belonging to the novel virus group, Salterprovirus. Virology 350:228–239.

65. Roux S, Solonenko NE, Dang VT, Poulos BT, Schwenck SM, Goldsmith DB, Coleman ML, Breitbart M, Sullivan MB. 2016. Towards quantitative viromics for both double-stranded and single-stranded DNA viruses. PeerJ 4:e2777.

66. Shakya M, Soucy SM, Zhaxybayeva O. 2017. Insights into origin and evolution of α-proteobacterial gene transfer agents. Virus Evol 3:vex036.

67. McDaniel LD, Young E, Delaney J, Ruhnau F, Ritchie KB, Paul JH. 2010. High frequency of horizontal gene transfer in the oceans. Science 330:50.

68. McDaniel LD, Young EC, Ritchie KB, Paul JH. 2012. Environmental factors influencing gene transfer agent (GTA) mediated transduction in the subtropical ocean. PLoS ONE 7:e43506.

69. Makhalanyane TP, Valverde A, Gunnigle E, Frossard A, Ramond J-B, Cowan DA. 2015. Microbial ecology of hot desert edaphic systems. FEMS Microbiol Rev 39:203–221.

70. Green MR, Sambrook J. 2016. Precipitation of DNA with Ethanol. Cold Spring Harb Protoc 2016:pdb.prot093377.

71. 2015. FastQC. https://qubeshub.org/resources/fastqc.

72. Bankevich A, Nurk S, Antipov D, Gurevich AA, Dvorkin M, Kulikov AS, Lesin VM, Nikolenko SI, Pham S, Prjibelski AD, Pyshkin AV, Sirotkin AV, Vyahhi N, Tesler G, Alekseyev MA, Pevzner PA. 2012. SPAdes: a new genome assembly algorithm and its applications to single-cell sequencing. J Comput Biol 19:455– 477.

73. Langmead B, Salzberg SL. 2012. Fast gapped-read alignment with Bowtie 2. Nat Methods 9:357–359.

74. Chen I-MA, Markowitz VM, Chu K, Palaniappan K, Szeto E, Pillay M, Ratner A, Huang J, Andersen E, Huntemann M, Varghese N, Hadjithomas M, Tennessen K, Nielsen T, Ivanova NN, Kyrpides NC. 2017. IMG/M: integrated genome and metagenome comparative data analysis system. Nucleic Acids Res 45:D507–D516.

75. Wagner GP, Kin K, Lynch VJ. 2012. Measurement of mRNA abundance using RNA-seq data: RPKM measure is inconsistent among samples. Theory Biosci 131:281–285.

76. Love MI, Huber W, Anders S. 2014. Moderated estimation of fold change and dispersion for RNA-seq data with DESeq2. Genome Biol 15:550.

77. Kanehisa M. 2017. Enzyme Annotation and Metabolic Reconstruction Using KEGG. Methods Mol Biol 1611:135–145.

78. Altschul SF, Koonin EV. 1998. Iterated profile searches with PSI-BLAST--a tool for discovery in protein databases. Trends Biochem Sci 23:444–447.

79. Marchler-Bauer A, Bo Y, Han L, He J, Lanczycki CJ, Lu S, Chitsaz F, Derbyshire MK, Geer RC, Gonzales NR, Gwadz M, Hurwitz DI, Lu F, Marchler GH, Song JS, Thanki N, Wang Z, Yamashita RA, Zhang D, Zheng C, Geer LY, Bryant SH. 2017. CDD/SPARCLE: functional classification of proteins via subfamily domain architectures. Nucleic Acids Res 45:D200–D203.

80. Katoh K, Rozewicki J, Yamada KD. 2019. MAFFT online service: multiple sequence alignment, interactive sequence choice and visualization. Brief Bioinform 20:1160–1166.

81. Huang Y, Niu B, Gao Y, Fu L, Li W. 2010. CD-HIT Suite: a web server for clustering and comparing biological sequences. Bioinformatics 26:680–682.

82. Gouveia-Oliveira R, Sackett PW, Pedersen AG. 2007. MaxAlign: maximizing usable data in an alignment. BMC Bioinformatics 8:312.

83. Cambuy DD, Coutinho FH, Dutilh BE. 2016. Contig annotation tool CAT robustly classifies assembled metagenomic contigs and long sequences. bioRxiv 072868.

84. Brister JR, Ako-adjei D, Bao Y, Blinkova O. 2015. NCBI Viral Genomes Resource. Nucleic Acids Res 43:D571–D577.

85. Huson DH, Mitra S, Ruscheweyh H-J, Weber N, Schuster SC. 2011. Integrative analysis of environmental sequences using MEGAN4. Genome Res 21:1552–1560.

86. Bolduc B, Jang HB, Doulcier G, You Z-Q, Roux S, Sullivan MB. 2017. vConTACT: an iVirus tool to classify double-stranded DNA viruses that infect Archaea and Bacteria. PeerJ 5:e3243.

87. Grazziotin AL, Koonin EV, Kristensen DM. 2017. Prokaryotic Virus Orthologous Groups (pVOGs): a resource for comparative genomics and protein family annotation. Nucleic Acids Res 45:D491–D498.

88. Johnson LS, Eddy SR, Portugaly E. 2010. Hidden Markov model speed heuristic and iterative HMM search procedure. BMC Bioinformatics 11:431.

89. Guy L, Nystedt B, Toft C, Zaremba-Niedzwiedzka K, Berglund EC, Granberg F, Näslund K, Eriksson A-S, Andersson SGE. 2013. A gene transfer agent and a dynamic repertoire of secretion systems hold the keys to the explosive radiation of the emerging pathogen Bartonella. PLoS Genet 9:e1003393.

