## Supplemental Data 2 for "With a pinch of salt: metagenomic insights into Namib Desert salt pan microbial mats and halites reveal functionally adapted and competitive communities"

**Supplementary Material**

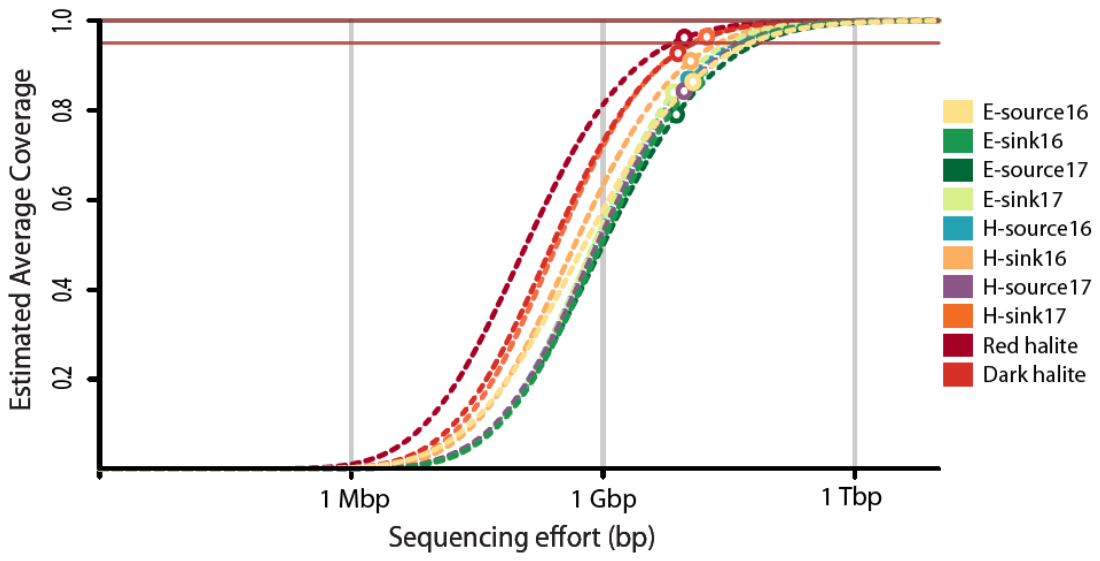

**Supplementary Figure 1. Estimation of the microbial community complexity and of the community coverage of the metagenomic datasets by Nonpareil.** The dark and light red horizontal lines indicate 100 and 95% coverage, respectively. The empty circles indicate the estimated coverage of each sample.

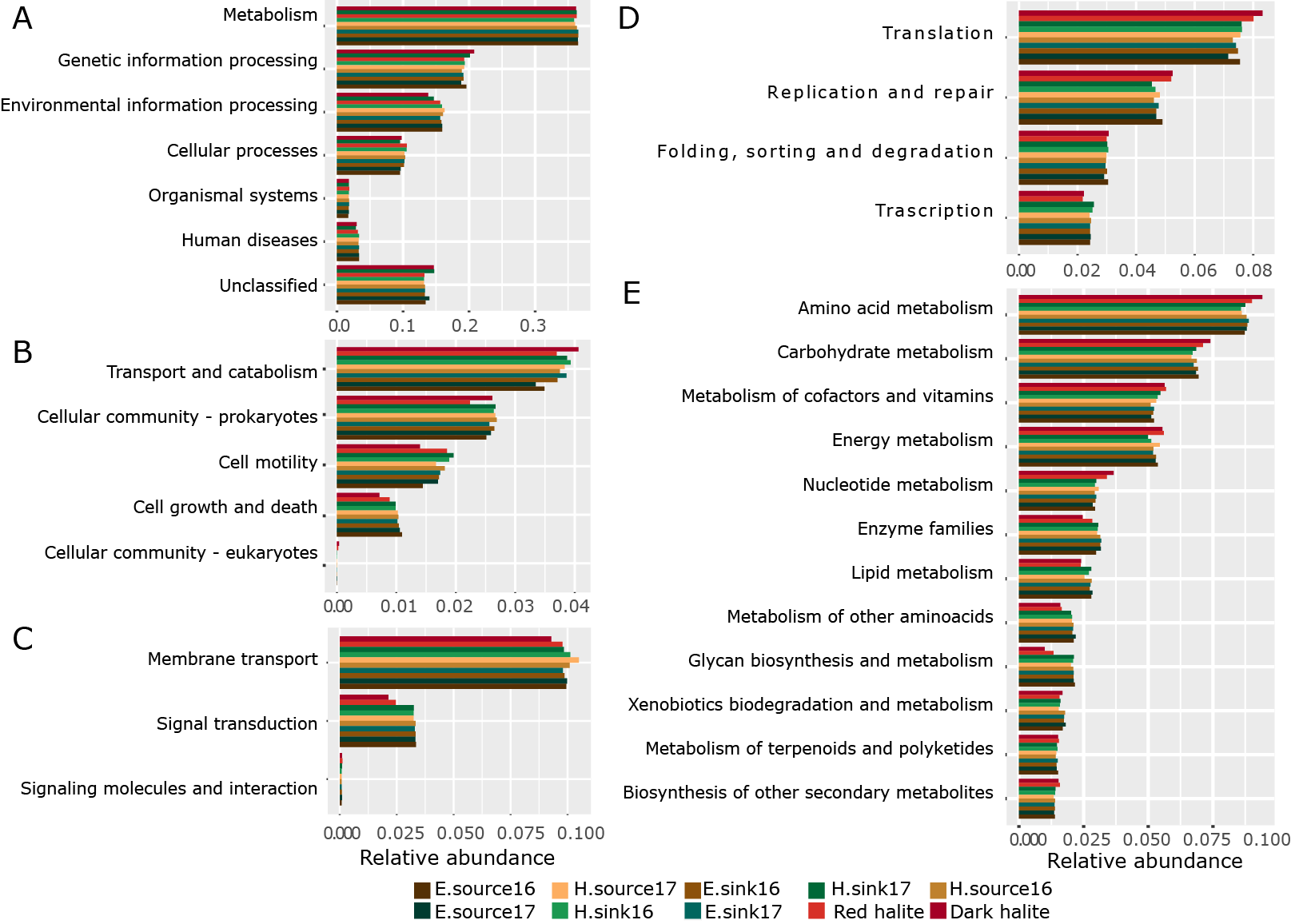

**Supplementary Figure 2. Relative abundance of metagenomic ORFs classified by the category or subcategory of their KEGG Orthology annotation. A.** Classification by KEGG categories. **B.** Classification within the category of Cellular Processes. **C.** Classification within the category of Environmental information processing. **D.** Classification within the category of Genetic information processing. **E.** Classification within the category of Metabolism.

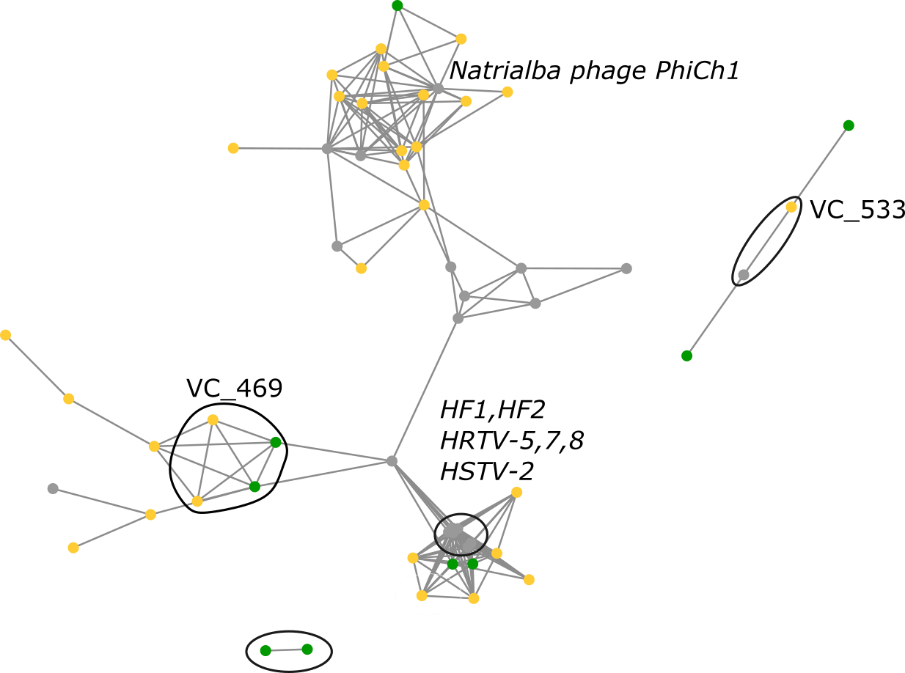

**Supplementary Figure 3. Comparison of halite viral sequences from the Namib and the Atacama Desert.** Genome-based protein sharing network of metagenomic halite viruses from the Namib Desert (yellow) and the Atacama Desert (green). Viral clusters (groups of sequences equivalent to viral genera) identified by the tool vContact2 are circled in black for clarity.

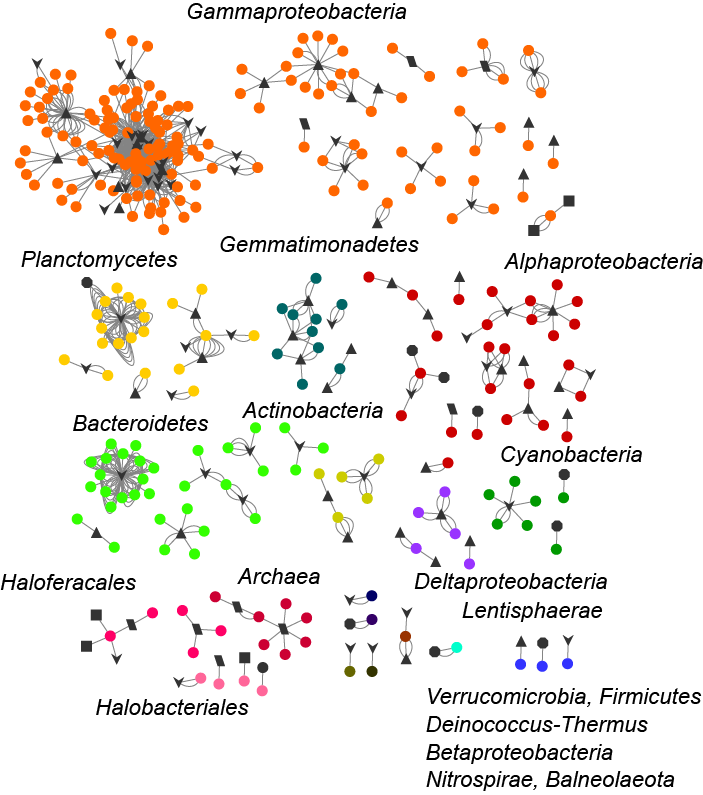

**Supplementary Figure 4. Map of virus-host interactions in the Namib Desert stream mat and halite microbial communities.** Circles depict host contigs, colored accordingly to their taxonomic assignment. Black symbols represent viral contigs, and the shape of the figure represents the source of the viral sequence. Lines represent a single spacer match.

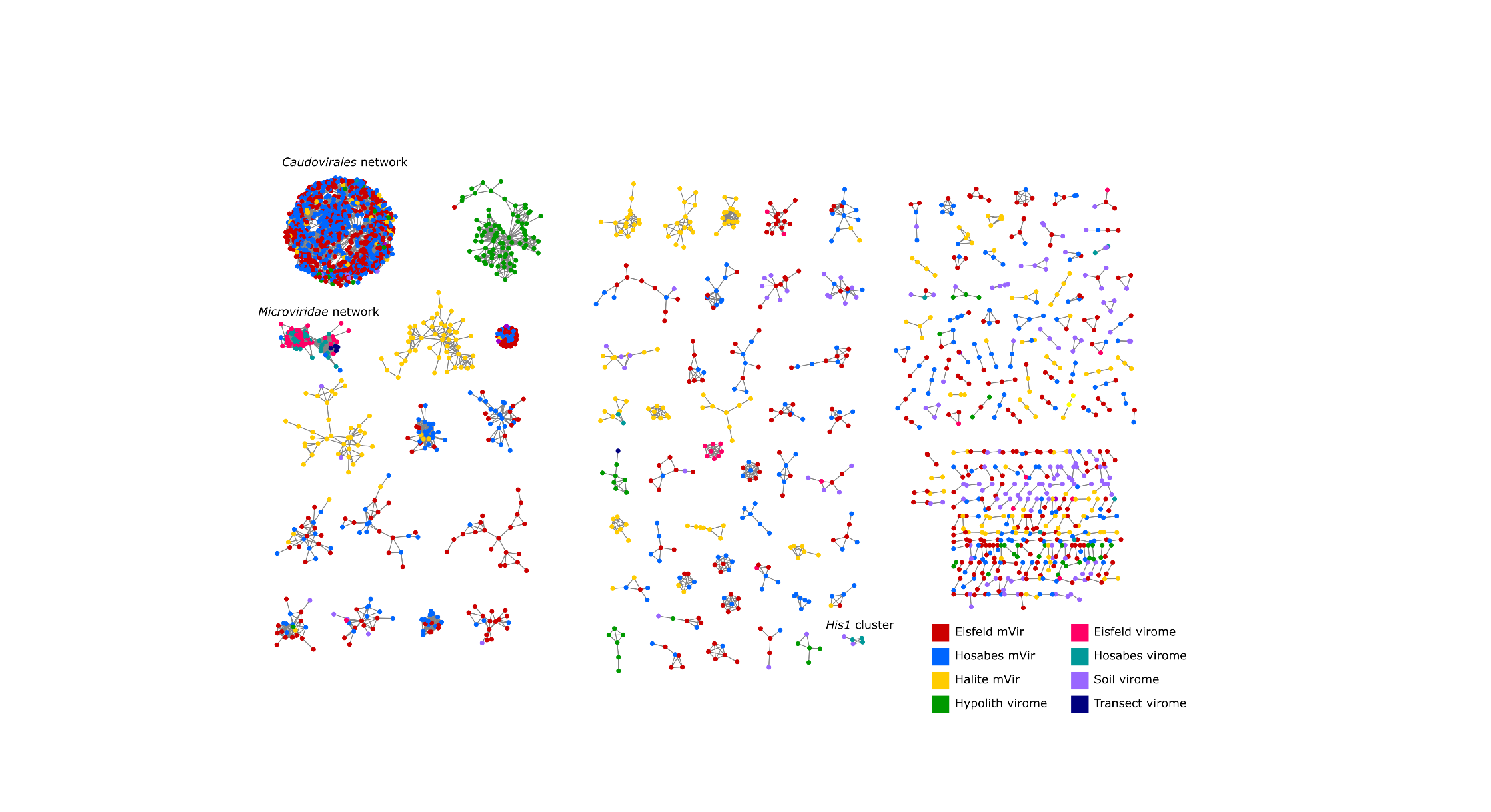

**Supplementary Figure 5. Genome-based protein-sharing network of viral sequences from the NamibVir and mVir datasets.** Each node represents a viral genome and edges represent statistically significant relationships between the protein profiles of those viral genomes.

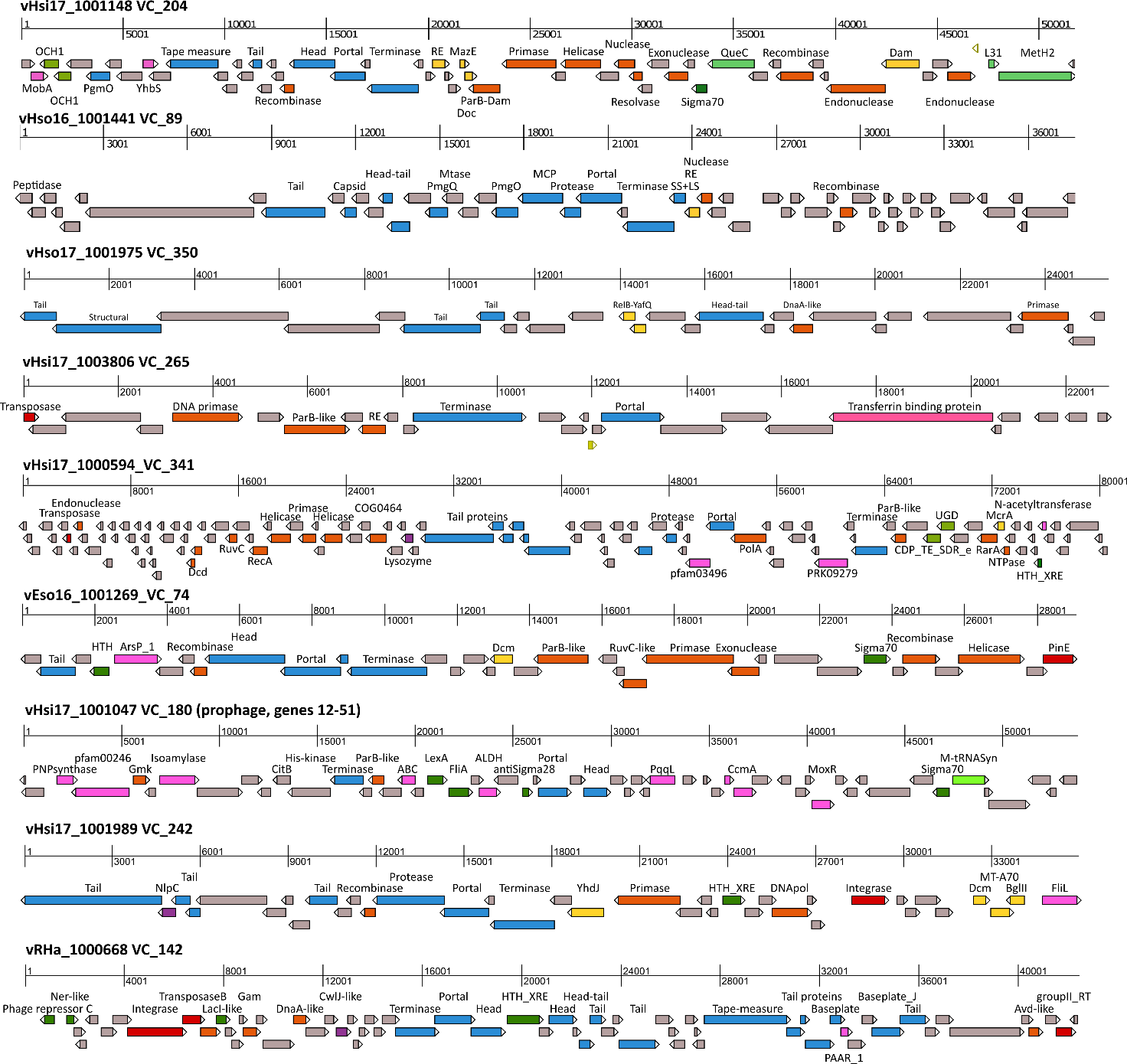

**Supplementary Figure 6. Genome organization of representative contigs from the largest mVir clusters.** Genes are colored according to their functional category: structural (blue), DNA metabolism (orange), lysis (purple), defense (yellow), transposases and integrases (red), transcription (dark green), translation (light green), metabolism (pink), unclassified (gray), tRNAs (yellow arrows).

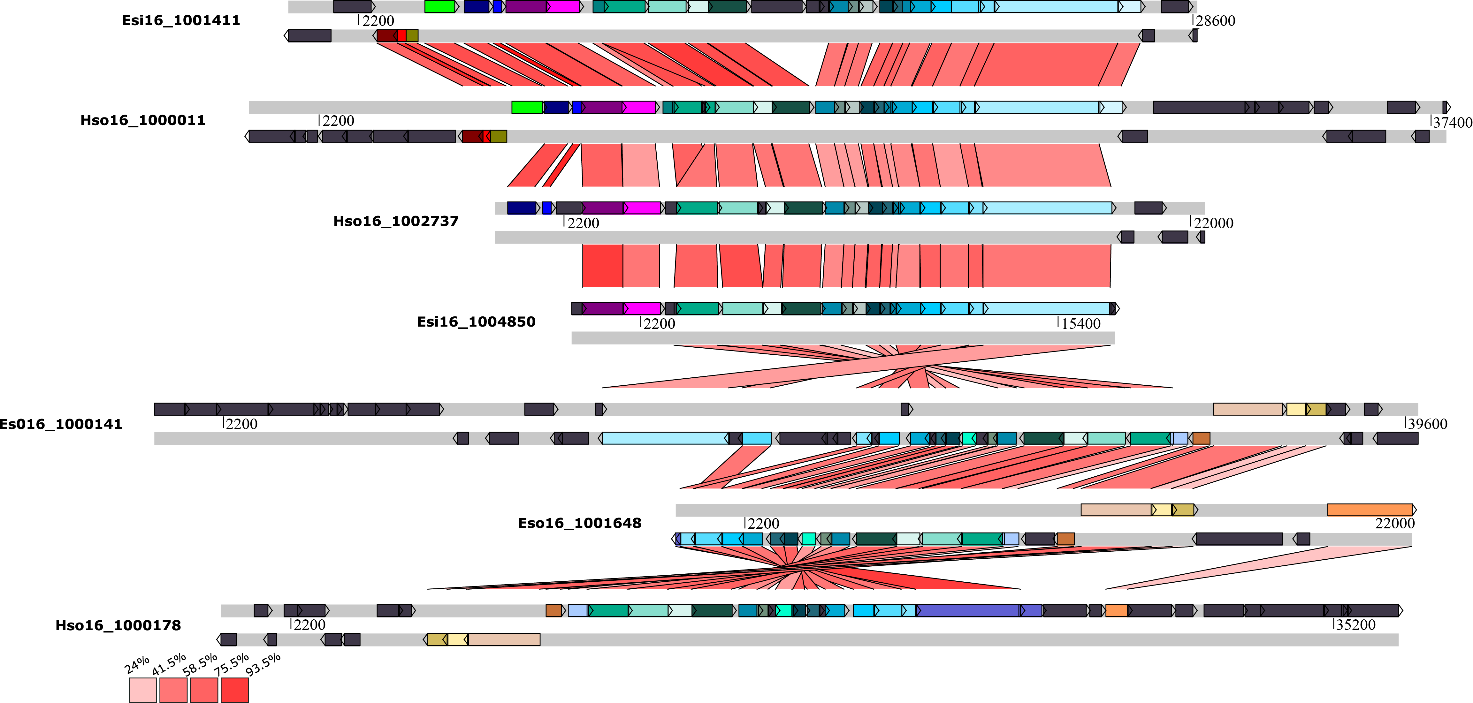

**Supplementary Figure 7. Genomic neighborhoods of the VC_69 contigs.** Flanking regions of 10 genes at both sides of each viral sequence were retrieved and used to calculate pairwise similarities between a pair of genomes (indicated as shades of red). Genes are depicted as arrows and homologs are color-coded for clarity, while genes without homology between a pair of genomes are shown in dark grey. Genes annotated as viral are shown in shades of light blue.

**Supplementary Table legends**

**Supplementary Table 1. Results of the sequencing, quality control and assembly of the Namib stream mat and halite metagenomes.**

**Supplementary Table 2 Taxonomic composition of the Namib stream mat microbial metagenomes at family level**

**Supplementary Table 3. Differential abundance of microbial families in the Eisfeld and Hosabes metagenomes**

**Supplementary Table 4. Differential abundance of microbial taxa in the source and sink of the stream mats**

**Supplementary Table 5. Taxonomic composition of the Namib halite metagenomes at family and genus level**

**Supplementary Table 6. Normalized matrix of KO terms per KEGG Module.** Eso – Eisfeld source; Esi – Eisfeld sink; Hso – Hosabes source; Hsi –Hosabes sink; Ha – halite.

**Supplementary Table 7. Raw counts of selected marker genes related to nutrient cycling and their distribution at family level**

**Supplementary Table 8. Identification of prokaryotic defence systems with PADLOC.**

**Supplementary Table 9. Predicted *cas* genes in the Namb Desert salt-pan metagenomes and subtype classification of Cas modules.**

**Supplementary Table 10. Relative abundance of the metagenomic *cas9* genes classified by their taxonomic affiliation**

**Supplementary Table 11. Viral sequences identified from the Namib stream mat and halite metagenomes**

**Supplementary Table 12. Gene-based genome network comparison of mVir, NamibVir and RefSeq viruses.**

**Supplementary Table 13. Overview of genome-to-cluster assignment by vContact 2.0**

**Supplementary Table 14. Overview of the viral clusters defined by vContact 2.0**

**Supplementary Table 15. Viral clusters taxonomy assignment**

**Supplementary Table 16. Gene-based genome sharing comparison of mVir and halite viruses from the Atacama Desert**

**Supplementary Table 17. Virus-host linkages identified through CRISPR spacer matches**

**Supplementary Table 18. Virus-host linkages identified through metagenomic contigs harboring prophages**

**Supplementary Table 19. Gene-based genome network comparison of unfiltered mVir and NamibVir datasets.**

**Supplementary Table 20. Functional annotation of mVir ORFs**

**Supplementary Table 21. Protein BLAST RcGTA and BaGTA genes to the mVir dataset**
